# Ready-to-load MHC-I Nanoparticles for High-throughput T cell Screening Studies

**DOI:** 10.1101/2024.12.17.628918

**Authors:** Hoang Anh T. Phan, Daniel Hwang, Michael C. Young, Shirley M. Sun, Dimitri S. Monos, Nikolaos G. Sgourakis

## Abstract

MHC-I proteins present epitopic peptides to CD8+ T cells to elicit multifaceted adaptive immune responses. The affinity and avidity of interactions between peptide-MHC molecules and T-cell receptors (TCR) are fundamental parameters that contribute to the induction of activated or anergic T cell states. Here, we present a loadable system, VLP-Open HLA, featuring a virus-like particle (VLP) that can accommodate up to 60 loadable HLA (HLA – human leukocyte antigen) molecules. HLA nanoparticles, pre-loaded with a placeholder ligand, allow efficient peptide exchange upon incubation with target peptides. We show that fluorescently tagged VLP-Open HLA particles can be used to stain antigen-specific CD8+ T cells, providing a screening tool for novel TCRs. Finally, we demonstrate that our system can induce activation of T cells in an antigen-specific manner. Our platform can be adapted to encompass multiple HLA allotypes and co-stimulatory molecules as mosaic nanoparticles, to enable a range of applications in experimental immunology.

## Introduction

Major histocompatibility complex class I (MHC-I) molecules play a central role in adaptive immunity of jawed vertebrates. MHC-I proteins comprise a polymorphic heavy chain (HC), an invariable light chain β_2_ microglobulin (β_2_m), and an epitopic peptide of 8-15 amino acid in length which enables immunosurveillance of the intracellular proteome^1,2^. MHC-I molecules are assembled in the endoplasmic reticulum (ER) and loaded with a pool of up to 10^5^ immunogenic peptides for trafficking to the cell surface^3^. In the human population, the highly polymorphic Human Leucocyte Antigen proteins (HLA; the MHC in humans comprising >30,000 allomorphs) present distinct repertoires of endogenous peptides, thereby ensuring species adaptability to emerging intracellular pathogens^4^. Specific engagement between an antigen-presenting cell and a CD8^+^ T cell is conferred by the intermediate affinity (micromolar range K_D_) interaction between a peptide-loaded MHC molecule (pMHC) and a clonotypic αβ T cell receptor (TCR), expressed on the T cell surface^5^. Robust T cell activation and proliferation towards a sustained immune response is dependent upon engagement of co-stimulatory receptors, adhesion molecules, and necessitates a sufficient TCR:pMHC interaction to enact CD45 phosphatase exclusion and ultimately TCR-CD3 complex triggering via distinct mechanisms^6^. Therefore, due to short half-life of the TCR:pMHC interaction, soluble monomeric pMHC-I molecules are insufficient to detect or to efficiently activate their antigen-specific T cells, and the immunology community has relied on multimerized MHC-I conjugates to identify cross-reactive T cells from polyclonal repertoires, and to monitor responses in various settings^7^. While several elegant approaches have been developed to generate libraries of MHC-I molecules encompassing hundreds of different epitopic peptides^8–12^, multimerized MHC-I molecules insufficiently stimulate T cells responses and are therefore limited to being used as probing reagents. There is thus a need for systems that can rapidly and accurately screen for epitopic peptides that can elicit specific T cell responses for immunological studies, and therapeutics development.

To overcome this challenge, Schneck and colleagues have developed nanoscale artificial antigen-presenting cells (aAPCs) for enrichment and expansion of antigen-specific T cells including iron-dextran paramagnetic nanoparticles (50-100 nm diameter), avidin-coated quantum dot nanocrystals (30-nm diameter), and larger size aAPCs (300-nm diameter), as well as magnetic nanoparticle-based for detection of antigen-specific T cells^13–16^. aAPCs can activate and rapidly expand antigen-specific T cells from melanoma patients^17,18^. pMHC-based aAPCs can expand T cells that are specific for cancer and autoimmune epitopes^19–21^. Other multimerization approaches for T cell activation include functionalization of Strep-Tactin (mutein of streptavidin) with anti-CD3 and anti-CD28 antibody Fab fragments (Expamers)^22^. Finally, CombiCells, a synthetic biology system enables combinatorial display of cell surface ligands, enabling studies of T-cell antigen sensitivity as well as modulation by co-stimulation or co-inhibition repceptors^23^. Recently, protein nanoparticles and cage-based protein assemblies have been gaining attention in the context of vaccine design^24,25^. Virus-like particles (VLPs) can offer multivalency and a symmetry that has been shown to be beneficial for antigen presentation in the context of vaccine development^26^. Built upon the 60-subunit nanocage based on 2-keto-3-deoxy-phosphogluconate (KDPG) aldolase developed by Baker *et al.*, Howarth, Townsend, and colleagues developed VLPs displaying the SARS-CoV-2 spike protein receptor-binding domain utilizing SpyCatcher/SpyTag chemistry^26–30^. The same group established the SpyCatcher003 scaffold that enabled efficient VLP conjugation and enhanced stability^26,30^. To functionalize the nanoparticles, SpyCatcher/SpyTag chemistry offers irreversible isopeptide bond formation via site-specific conjugation between a lysine residue on the SpyCatcher protein and an aspartic acid on its SpyTag protein partner.^31^

In this study, we aim to develop a universal, loadable VLP-based nanoparticle platform to screen for peptide/MHC-I restricted T cell responses. Several methods have been proposed for generating loadable pMHC molecules, including UV irradiation of a photocleavable conditional ligand, disulfide bond direct stabilization of the MHC-I peptide-binding groove, or elevated temperature treatment^8,32–36^. We have previously established a high-throughput method for chaperone-mediated exchange of a placeholder peptide for high-affinity peptides of choice^9^. Recently, we used a structure-guided approach to develop conformationally stabilized, open or ready-to-load MHC-I molecules^37^. Here, we leverage our open MHC-I system, the enhanced avidity of VLPs, and SpyCatcher/SpyTag-mediated multimerization, to develop a loadable open MHC nanoparticle platform that is compatible with different HLA allomorphs (VLP-Open HLA). Nanoparticles conjugated with open MHC molecules allow efficient peptide exchange *in vitro*, resulting in a stable pHLA display platform. As a proof of concept, we demonstrate that our VLP-Open HLA (Open HLA-A*02:01/NY-ESO-1) can be fluorescently tagged to stain antigen-specific NY-ESO-1 TCR (1G4) CD8+ T cells, and can specifically activate T cells in a detectable manner *in vitro*. Our platform provides a minimalistic protein-based system to systematically screen and evaluate T cell responses against a wide range of peptide/HLA combinations.

## Results

### Design and Characterization of VLP-Open HLA Conjugates

To generate VLP-Open HLA nanoparticles, we displayed SpyTagged open MHC on VLP-SpyCatcher-mi3. First, the heavy chain ectodomain of the open HLA-A*02:01 was fused to SpyTag003 via a Gly-rich linker (see Supporting Information, **Table S2**). Our engineered open HLA-A*02:01 features an interchain disulfide bond between the heavy (G120C) and light chain, β_2_m (H31C) (**Figures 1A-1B**), as previously described^37^. Compared to native molecules, the engineered open MHC I shows enhanced stability and rapid loading of exogenous peptides through the stabilization of a peptide-receptive conformation. This provides us with a ready-to-load MHC I system suitable for probing T cell activity and low-affinity ligand screening^37^. We refolded SpyTag open HLA-A*02:01 with the high-affinity peptides NY-ESO-1^C9V^ (SLLMWITQV), the mutated decoy peptide NY-ESO-1^W5A^ (SLLMAITQV), and a moderate-affinity placeholder peptide gTAX (LFGYPVYV) (**Figure 1C**). We find that adding the SpyTag does not compromise the thermal stability of the protein, evident by the high T_m_ values of the SpyTagged open HLA-A*02:01 refolded with different peptides between 51.8 °C and 58.9 °C using Differential Scanning Fluorimetry (DSF) (Supporting Information**, Figure S2**).

**Figure 1.**
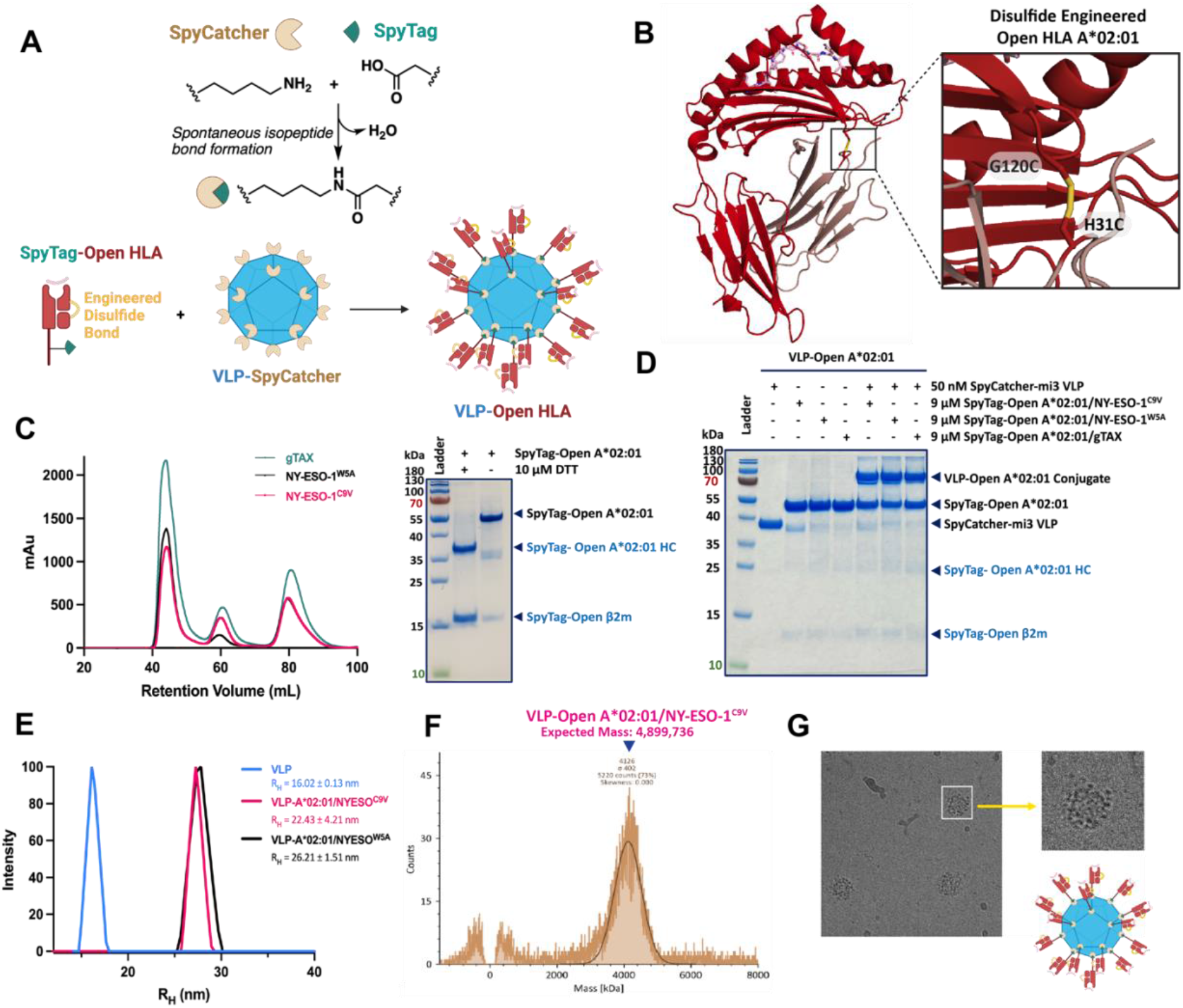
Design, production, and biophysical characterization of VLP-Open HLA system. **(A)** Schematic of the design: a SpyTag Open HLA molecule is conjugated to VLP-SpyCatcher to form VLP-Open HLA molecule. **(B)** Model of Open HLA-A*02:01 with an engineered disulfide bond as previously described by our group^37^. **(C)** SEC traces of the SpyTagged G120C/H31C “Open HLA-A*02:01” molecules refolded with NY-ESO-1^CV9^ (pink), NY-ESO-1^W5A^ (black), NY-ESO-1^gTAX^ (green). Representative SDS-PAGE analysis of SpyTagged Open A*02:01. **(D)** Optimized conjugation reaction between SpyCatcher-mi3 VLP and Open A*02:01 molecules refolded with different peptides as validated with SDS PAGE gel. The gel showed pre- and post-conjugation products. In SDS-PAGE, the 60-mer complex broke apart and was observed as a monomer band at ∼84 kDa. **(E)** Dynamic Light Scattering (DLS) scans of VLP-SpyCatcher-mi3 only, VLP-Open A*02:01/NY-ESO-1^C9V^, and VLP-Open A*02:01/NY-ESO-1^W5A^. **(F)** Representative mass photometry data for VLP-Open A*02:01/NY-ESO-1^C9V^, showing successful conjugation to most of the 60 sites of VLP. **(G)** Example from a cryo-electron micrograph of VLP-Open A*02:01/NY-ESO-1^C9V^.

We chose the protein-based nanoparticle platform virus-like particle (VLP) due to its versatility, ease of assembly, stability, and geometry ideal for antigen display.^26,38^ Specifically, we focused on the SpyCatcher-mi3 scaffold (with 60 binding sites on VLP) that enabled efficient VLP conjugation and enhanced stability, and can be expressed in *E.coli* (see Supporting Information, **Figure S1** for purification and production information)^38,30,26^. For the conjugation reaction, we optimized the molar ratio of SpyCatcher-mi3 and SpyTag open HLA-A*02:01 (see Supporting Information, **Table S3**). Our results suggested that a ratio of 1:180, with final concentrations such as 50 nM SpyCatcher-mi3 particle and 9 μM SpyTag Open HLA resulted in an optimal conjugation efficiency. We then conjugated SpyTag Open HLA-A*02:01 refolded with either NY-ESO-1^C9V^, NY-ESO-1^W5A^, or gTAX peptide; a band shift on SDS-PAGE gel confirmed the conjugation via covalent bond (**Figure 1D**). Any aggregate and unconjugated Open HLA molecule was removed using a high molecular weight cutoff (100 kDa MWCO) centrifugal spin column prior to characterization and subsequent experiments.

The robust conjugation reaction via SpyCatcher-SpyTag chemistry resulted in the formation of VLP-Open HLA protein nanoparticle, as further confirmed by dynamic light scattering (DLS). The size of our VLP only particle (R_H_ = 16.02 ± 0.12 nm) was consistent with the literature (R_H_ = 18.6 ± 4.8 nm from Tan *et al.,* 2021) (**Figure 1E**)^30^. The increase in particle size upon conjugation with Open HLA-A*02:01/NY-ESO-1^C9V^ or Open HLA-A*02:01/NY-ESO-1^W5A^ was consistent with the increase in size when conjugated of the size HLA molecule (∼47 kDa and 411 amino acids). We further characterized the particles using mass photometry, and the masses observed were consistent with the expected values for the unconjugated and conjugated particles (**Figure 1F** and Supporting Information, **Figure S4**). VLP-Open A*02:01/NY-ESO-1^C9V^ showed a mass distribution for the conjugated particle at 4126 ± 402 kDa, indicating an average occupancy of approximately 46 out of 60 sites on the VLP by pHLA molecules (**Figure 1F**). Cryo-electron microscopy (cryo-EM) further confirmed the presence of a cage-like structural arrangement (**Figure 1G**). Together, these studies support robust presentation of open MHC I molecules by VLP nanoparticles using *in vitro* assembly from recombinant protein expression in *E. coli*.

### VLP-Open HLA molecules can capture exogenous peptides *in vitro*

We previously established that our engineered open MHC-I design enables enhanced peptide exchange of placeholder peptides across polymorphic HLA molecules^37^. We next examined whether Open HLA (Open HLA-A*02:01) conjugated to VLPs retain their enhanced peptide exchange properties, to evaluate its potential as a screening system for immunogenic peptides. Open HLA-A*02:01 was refolded with the placeholder peptide, gTAX, prior to conjugation on the VLP scaffold. Via fluorescence polarization (FP), we observed rapid peptide exchange of gTAX for the labelled peptide _TAMRA_TAX9 (TAMRA-KLFGYPVYV) on VLP-Open HLA (**Figures 2A-2B**). The association of fluorescently labeled peptides was consistent across independent replicates of VLP sample preparation (Supporting Information, **Figure S5**). Peptide loading was dependent on the nanoparticle concentration, showing an overall similar profile as observed in our previous study using soluble MHC I molecules. We observed complete peptide exchange within 4 hours at room temperature, compared to minimal exchange using wild-type HLA-A*02:01^37^. Therefore, our VLP-Open HLA-A*02:01/gTAX design can promote efficient peptide exchange *in vitro*.

**Figure 2.**
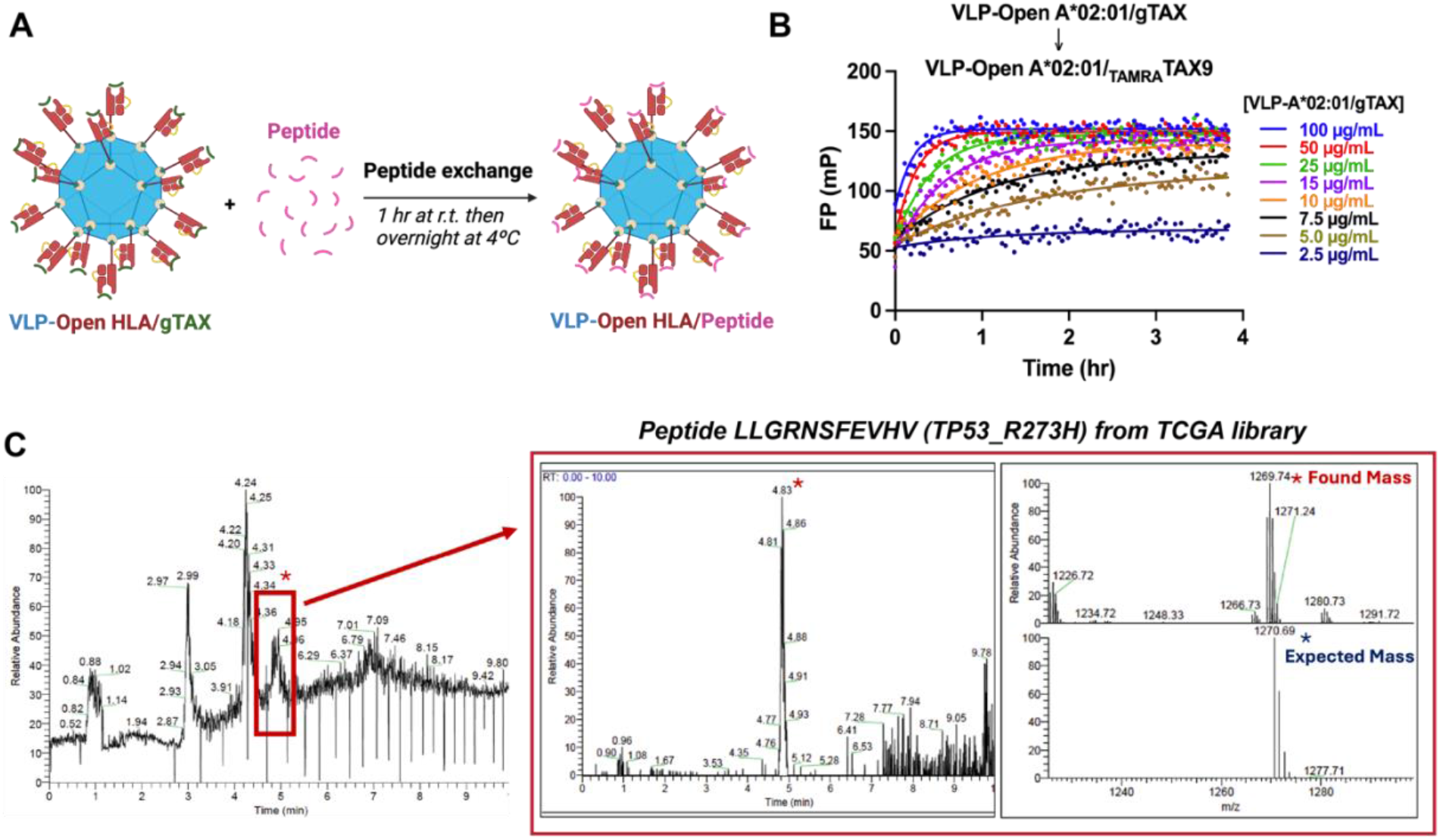
VLP-Open HLA can capture and display exogenous peptides. (A) Principle of peptide exchange on VLP-Open HLA system. Peptide exchange can take place at room temperature for 1 hour at room temperature then overnight at 4°C (procedure used for subsequent immunological assays). (B) Loading and Binding of _TAMRA_TAX9 by VLP-Open A*02:01/gTAX monitored at room temperature. Peptide exchange was measured by fluorescence polarization (mP) of 40nM _TAMRA_TAX9 as a function of the VLP-Open A*02:01/gTAX concentrations. Individual traces were fit to an exponential association model. The means from each condition’s triplicates were plotted. (C) Representative LC-MS Mass Spectrometry analysis of peptide bound to VLP-Open A*02:01; the peptide is from TCGA peptide library. VLP-Open A*02:01/gTAX can load and capture different antigenic peptides.

Furthermore, we conducted mass spectrometry experiment to demonstrate that our platform can be used to identify potentially immunogenic peptides from biological samples. As a proof of concept, VLP-Open HLA-A*02:01/gTAX was exchanged with either individual peptides, or a set of 50 candidate high-affinity peptides derived from the Cancer Genome Atlas (TCGA), previously validated by our group^37^ (for peptide information, see Supporting Information, **Table S4**). Excess peptides were washed using spin columns to retain bound peptides; a negative control with peptide only (no VLP-Open HLA particle) and a negative control with VLP-Open HLA particle in the presence of an irrelevant peptide were included to ensure that our method efficiently washed out the excess peptide (Supporting Information, **Figures S6-S7**). Captured peptides were identified using liquid chromatography-mass spectrometry (LC-MS). As expected, we found that the predicted high-affinity peptides were retained on the VLP-Open HLA nanoparticle, and readily identified in our data (**Figure 2C)**. Additional mass spectrometry data from incubation of VLP-Open HLA-A*02:01/gTAX with individual or mixture of peptides from the TCGA epitope library are shown in the Supporting Information, using two complementary methods – LC-MS (Supporting Information, **Figures S8-S11)** and matrix-assisted laser desorption/ionization time-of-flight mass spectrometer (MALDI-MS) **(Figure S13)**.

### Fluorescently Labeled Open HLA-VLPs can detect antigen-specific T Cells

VLP-SpyCatcher-mi3 can be fluorescently labeled by reaction with NHS-fluorescein prior to conjugation with molar excess amount of Open HLA molecule without compromising its pHLA conjugation capacity (**Figure 3A**). From SDS-PAGE analysis, under UV radiation and prior to Coomassie staining, we can observe that post-labeled sample showed a green band corresponding to SpyCatcher-mi3 (VLP is shown as monomer in reducing condition on SDS-PAGE gel) (**Figure 3B**). Mass spectrometry data (MALDI-MS) confirmed that the VLP monomer was effectively conjugated with fluorescein, evident by the mass shift corresponding to 3-5 lysine residues labeled (Supporting Information, **Figure S14**).

**Figure 3.**
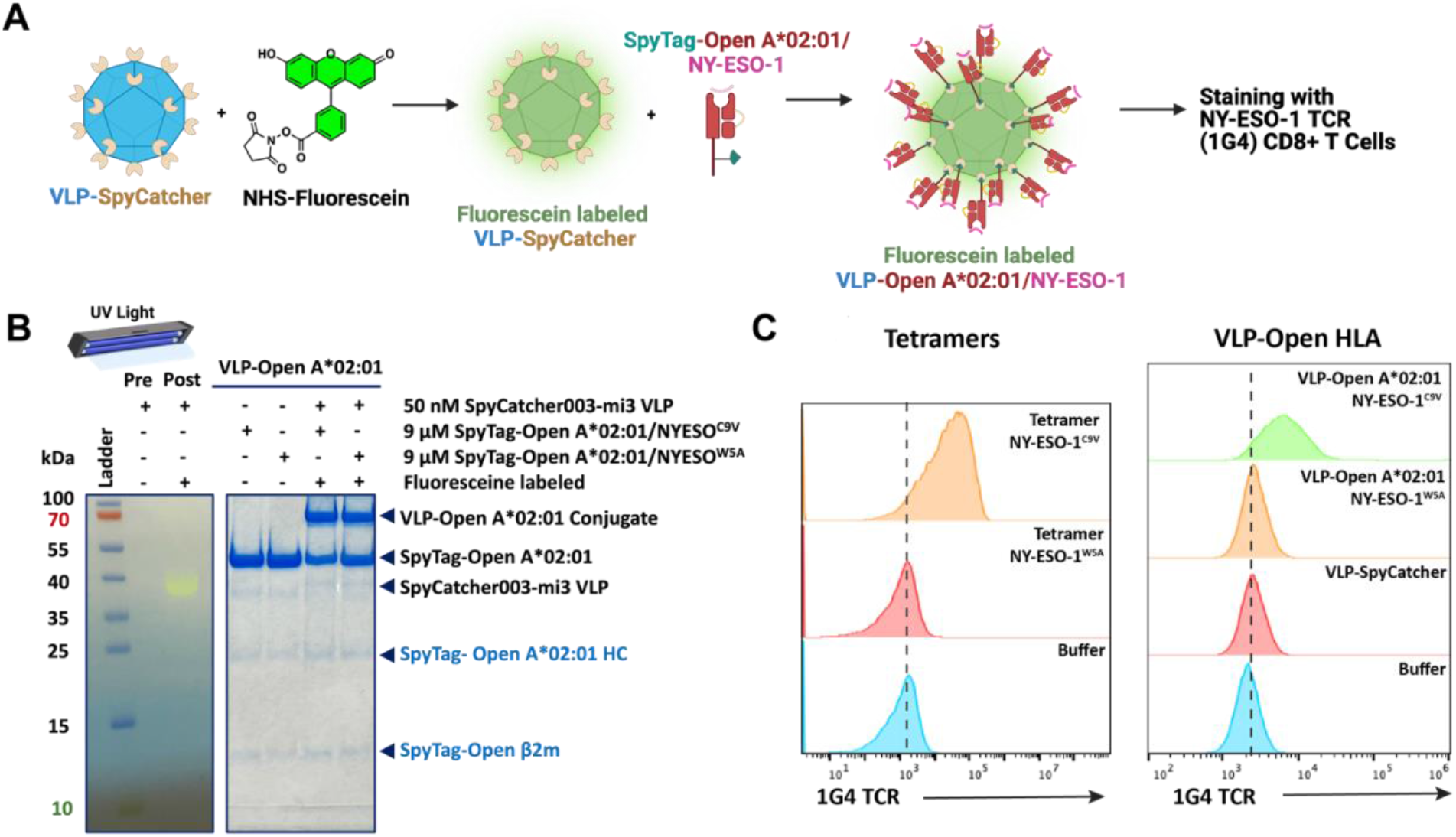
The VLP-Open HLA system can detect antigen-specific T cells. (**A**) Labeling Scheme for VLP Open-HLA. **(B)** SDS-PAGE analysis of SpyCatcher-mi3 VLP pre- and post-labeling with fluorescein; without gel staining and under UV-light, the post-labeling sample showed a green band at the expected molecular weight of SpyCatcher-mi3, indicative of a fluorescently labeled VLP complex. This was then successfully conjugated to Open A*02:01/ NY-ESO-1^C9V^ or Open A*02:01/NY-ESO-1^W5A^ for T cell staining experiment. (**C**) 1G4 CD8+ T cell stained with tetramers (controls; *left panel*) and with VLP-Open-HLA refolded with NY-ESO-1^C9V^ or NY-ESO-1^W5A^ (*right panel*). Gating strategy is shown in the Supporting Information, **Figure S15**.

The labeled SpyCatcher-VLP was then successfully conjugated to Open HLA-A*02:01 refolded with NY-ESO-1^C9V^ or NY-ESO-1^W5A^ using our established conjugation protocol (**Figure 3B**). We then explored whether our fluorescently labeled VLP-Open HLA-A*02:01/NY-ESO-1^C9V^ could be used to detect antigen-specific CD8+ T cells, using primary human T cells expressing the 1G4 TCR specific for the cancer testis antigen NY-ESO-1 (**Figure 3C**)^39^. We used tetramerized Open HLA-A*02:01/NY-ESO-1^C9V^ molecules to confirm antigen-specific staining of our engineered 1G4 T cells (positive control), while tetramer of Open HLA-A*02:01/NY-ESO-1^W5A^, which contains a key amino acid substitution to NY-ESO-1^C9V^ necessary for high-affinity interactions with the 1G4 TCR, showed low background staining levels (negative control) (**Figure 3C**; left panel). We found that VLP-Open HLA-A*02:01/NY-ESO-1^C9V^ showed a clear median fluorescence intensity (MFI) shift, allowing efficient staining and detection of NY-ESO-1 TCR (1G4) CD8+ T cells (**Figure 3C**; right panel).

### Open HLA-VLP nanoparticles elicit antigen-specific T cell activation *in vitro*

To assess whether our VLP-Open HLA-I platform could elicit antigen-specific activation and proliferation of T cells, we conjugated Open HLA-A*02:01 refolded with NY-ESO-1^C9V^ to VLPs and stimulated CD8+ 1G4 TCR-T cells *in vitro* (**Figure 4A**). We found that VLP-Open HLA-A*02:01 at very low concentrations could elicit antigen-specific proliferation, as assessed by limiting dilution analysis of CellTraceViolet labeled 1G4 TCR-T cells. At the lowest tested concentration (0.39 µg/mL; VLP complex is about 4.9 MDa), VLP-Open HLA-A*02:01 induced proliferation at equivalent levels as commercial Human T-Activator anti-CD3/anti-CD28 beads (**Figure 4B, Figure S17**). Proliferation at this concentration was antigen-specific, as stimulation with VLPs presenting NY-ESO-1^W5A^ only triggered low levels of proliferation. Quantification of T cell activation markers showed similar trends, with antigen-specific induction of 4-1BB expression up to 1.56 μg/mL (**Figure 4C**; Supporting Information, Figure **S17**). Consistent with these trends, activation with VLP-Open HLA-A*02:01 particles elicited antigen-specific production of IFN-γ as measured by enzyme-linked immunosorbent assay (ELISA) (Supporting Information, **Figure S19**). Notably, at higher concentrations of VLP-Open HLA-I, HLA-I loaded with the low affinity NY-ESO-1^W5A^ peptide could elicit CD8+ 1G4 T cell activation, likely owing to the high avidity provided by the VLP-Open HLA-I platform.

**Figure 4.**
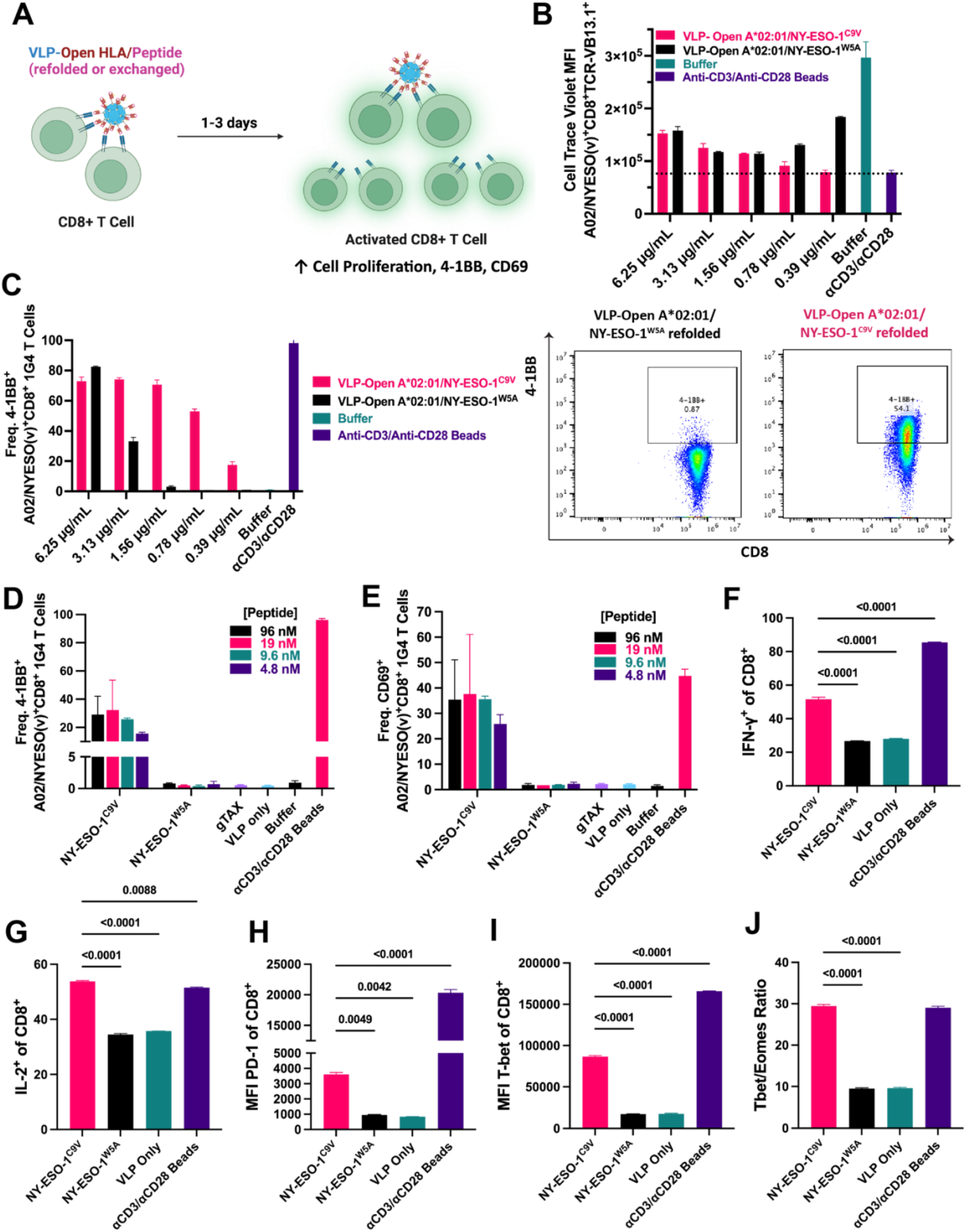
VLPs elicit antigen-specific activation of CD8+ T cells *in vitro*. **(A)** T-cell activation with Open HLA-VLP nanoparticles. **(B)** Exploring the Optimal Working Concentration Range of VLP-Open HLA-A*02:01/NY-ESO-1^C9V^ for Specific T cell Activation in a Cognate Antigen Dependent Manner. 1G4 CD8+ T cells were stimulated with different concentrations of VLP-Open HLA-A*02:01/NY-ESO-1^C9V^, VLP-Open HLA-A*02:01/NY-ESO-1^W5A^, media only (buffer negative control), or anti-CD3/anti-CD28 beads (positive control) for 3 days before flow cytometry analysis. Median fluorescence intensity (MFI) of Cell Trace Violet of CD8+ Live Cell population (lower intensity signals more population of proliferated cells). Average of 2 replicates is shown. **(C)** Frequency of 4-1BB+ cells in different conditions and representative flow cytometry data of 4-1BB+ CD8+ cell populations. **(D-E)** VLP-Open HLA-A*02:01/gTAX exchanged with NY-ESO-1^C9V^ can trigger T cell activation in a cognate antigen dependent manner. 1G4 CD8+ T cells were stimulated with VLP-Open HLA-A*02:01/gTAX exchanged with different concentrations of NY-ESO-1^C9V^ or NY-ESO-1^W5A^, VLP-HLA-A*02:01/gTAX only (no exchange), VLP only, media only (buffer negative control), or anti-CD3/anti-CD28 beads (positive control) for 3 days before flow cytometry analysis. Average of at least two replicates is shown. (**D)** Frequency of 4-1BB^+^ and (**E)** CD69^+^ cells among CD8^+^ cells. (**F-J)** Quantification of different markers of T cell activation/differentiation. 1G4 T cells were stimulated with VLP-Open HLA-A*02:01/NY-ESO-1^C9V^, VLP-Open HLA-A*02:01/NY-ESO-1^W5A^, VLP only (negative control), or anti-CD3/anti-CD28 beads. VLP were added at a concentration 0.78 μg/mL and incubated for 3 days. (**F-G)** After 3 days of activation, cells were stimulated to produce cytokines by addition of a cocktail of PMA/Ionomycin and brefeldin A: (**F)** Frequency of IFN-g^+^ and (**G)** IL-2^+^ cells among CD8+ T cells. (**H-J)** Expression of markers in cells following 3 days activation (without further PMA/Ionomycin stimulation): (**H)** MFI of PD-1, and (**I)** T-bet. (**J)** T-bet/Eomes ratio. Average of 2 replicates is shown, and significance was determined by One-way ANOVA followed by multiple comparisons testing.

We next turned our attention to testing whether a placeholder peptide loaded on VLP-Open HLA-A*02:01 could be exchanged for an antigen of interest, as this would enable a single stock of VLPs to be used for different HLA-A*02:01 binding peptides. We prepared VLP-Open HLA-A*02:01 loaded with a placeholder gTAX peptide and incubated them with up to 0.24 μM of NY-ESO-1^C9V^ and NY-ESO-1^W5A^ peptides (a concentration which would theoretically saturate all 60 binding sites; 1:60 VLP-HLA:Peptide molar ratio). Peptides were not washed out before use in these T cell activation assays. To control for any effects caused by excess peptide, peptides of the same concentrations (without VLP) were also prepared as peptide controls for the T cell activation assays (Supporting Information, **Figure S18**). Stimulation with VLP-Open HLA-A*02:01/gTAX exchanged with NY-ESO-1^C9V^ induced expression of 4-1BB and CD69 and production of IFN-γ from 1G4 T cells, while VLPs exchanged with NY-ESO-1^W5A^, or loaded with gTAX peptide, or peptides alone did not (**Figures 4D-E**; Supporting Information, **Figures S18 & S20**). The VLP-Open HLA-I system thus enables exchange of a placeholder peptide for a target peptide via simple incubation, without necessity for a wash step.

Lastly, we sought to evaluate common markers used for assessing T cell activation/differentiation following VLP stimulation. We first performed intracellular cytokine staining and found that VLP-Open HLA-A*02:01/NY-ESO-1^C9V^ stimulation increased the frequency of IFN-γ and IL-2 producing 1G4 T cells (**Figures 4F-G**). Similarly, activation/exhaustion marker PD-1 was upregulated in VLP-open HLA-A*02:01/NY-ESO-1^C9V^ stimulated cells, but not in VLP-Open HLA-A*02:01/NY-ESO-1^W5A^ or VLP only treated 1G4 T cells (**Figure 4H**). Lastly, we stained for T cell differentiation markers T-bet and Eomes and found increased expression of these transcription factors in cells stimulated with VLP-Open HLA-A*02:01/NY-ESO-1^C9V^ or with anti-CD3/anti-CD28 beads (**Figures 4I-J**; Supporting Information, **Figure S21**). The ratio of T-bet to Eomes expression was similar, indicating that VLP-Open A*02:01/NY-ESO-1^C9V^ drove effector T cell differentiation in a manner similar to that induced by anti-CD3/anti-CD28 beads. Together, these data suggest that VLP-Open HLA stimulation generates TCR signals similar to natural stimulation, eliciting measurable responses across a variety of different markers.

## Discussion

In the present work, we have described and developed the VLP-Open HLA, a loadable system featuring a virus-like particle protein cage and our previously described engineered open HLA proteins^40^. Utilizing SpyCatcher/SpyTag chemistry, we successfully produced and characterized our VLP-Open HLA particles. We demonstrated that these nanoparticles can be pre-loaded with a placeholder ligand and be efficiently exchanged with candidate peptides, showing its potential utility as a screening tool of peptide mixtures from biological samples. We further showed that fluorescently tagged VLP-Open HLA particles can be used to stain antigen-specific CD8+ T cells, providing a tool to evaluate the cross-reactivity of TCRs in different research and clinical settings. With a 60-mer dodecahedral geometry, our VLP-Open HLA is armed with both affinity and avidity properties for efficient engagements between peptide-MHC molecules and T-cell receptors, as being shown by its ability to not only recognize, but also to activate CD8+ T cells in an antigen specific manner.

Our platform can be adapted to present a wide range of patient-derived HLA I and HLA II allotypes and neoantigens as mosaic nanoparticles, to enable emerging cancer immunotherapy applications such as personalized T cell vaccines and T cell-based therapies^41^, while serving as a promising screening tool for patient specific immunotherapies. Specifically, beyond a screening and research tool, we envision that our VLP-Open HLA platform can be leveraged to make a loadable, multivalent system presenting MHC I molecules of different allotypes and antigen specificities for personalized polyclonal anti-tumor response. For instance, the VLP-Open HLA system can be conjugated with recombinant human leukocyte antigen (HLA) proteins expressed in a given patient, loaded with a set of candidate neoantigens to elicit polyclonal T cells specific to the patient’s tumor-specific HLA genotype and neoantigens. This can enable development of personalized T cell vaccines, autologous T cell-based therapies like tumor infiltrating lymphocytes (TILs), and screening tools for patient tumor specific synthetic immunoreceptors such as in Single-chain Fragment variable (ScFv) antibodies, Chimeric Antigen Receptors (CARs), or Bispecific T-cell Engagers (BiTEs). Overall, this manuscript aims to describe the initial design of the VLP-Open HLA platform that would lay the foundation for these promising future applications.

Moreover, the avidity demonstrated by the VLP-Open HLA-I platform with all sites conjugated to Open HLA-I molecules opens the door for future development of mosaic particles with co-stimulatory molecules co-conjugated with antigen-loaded HLA-I for many promising *in vitro* and *in vivo* applications. The size of VLP-Open HLA-I particle (∼30 nm) would enable widespread dissemination of VLPs *in vivo*, as they are sufficiently small to drain efficiently from the site of immunization directly to the lymphoid organs^42^. Therefore, it would be anticipated that an adequate dosage of VLP-Open HLA-I would be widely distributed to most populations of T cells throughout the body. It is then possible to envision conjugation of either positive or negative co-stimulatory molecules on the VLP surface, to drive robust T cell activation, survival, and expansion in cancer therapies^43^, or to direct anergy and exhaustion in autoimmunity diseases^44^. Such applications may have milder side effects, as VLP-Open HLA-I targeting may enable lower dosages to achieve desirable antigen-specific effects.

From both the oncological and autoimmune perspectives, the ability to load VLP-Open HLA-I particles with specific peptide cocktails are of significant interest, as this enables simultaneous stimulation or suppression of multiple antigen-specific T cell populations, potentially increasing the probability of sustained therapeutic efficacy. In conclusion, our platform provides a versatile, modular, and loadable system for influencing T-cell antigen-specific responses with a plethora of both *in vitro* and *in vivo* applications.

## Methods

### Expression & Purification of SpyTag Open HLA-A*02:01

The heavy chain of Open A*02:01 (G120C) ectodomain was C-terminally tagged with SpyTag003 (RGVPHIVMVDAYKRYK). This heavy chain was refolded with β2m (H31C) in the presence of excess peptide NY-ESO-1^C9V^ (SLLMWITQV), NY-ESO-1^W5A^ (SLLMAITQV; used negative control), or gTAX (LFGYPVYV; placeholder peptide). A Gly-rich linker (GGSGGGSGG) was included between the isopeptide fusion tag (SpyTag003) and the α3 domain of the heavy chain, to prevent interference between the tag function and the heavy chain and other undesired interactions. Plasmids encoding the luminal domain of the C-terminally linked SpyTag003 Open A*02:01 (G120C) and open human β2m (H31C) were transformed into *Escherichia coli* BL21(DE3) cells using the pET22b(+). The specific protein sequences are described in the Supporting Information, **Table S2**. The cells were grown and harvested in the Luria-Broth (LB) medium, and inclusion bodies were pelleted and purified as previously described.^37^ The ternary pMHC-I complexes were generated by an *in vitro* refolding, by slowly diluting a 100 mg of Open HLA-A*02:01-SpyTag003 and open β2m at a 1:3 molar ratio over 4 hours in 500 mL of refolding buffer (0.4 M L-Arginine HCl, 100 mM Tris, 2 mM EDTA, 5 mM reduced L-glutathione, 0.5 mM oxidized L-glutathione, pH 8.0) supplemented with 5 mg of the peptide. The refolding reaction proceeded for 4 days, and the proteins were then dialyzed into a buffer containing 25 mM Tris, 150 mM NaCl, pH 8.0 (TBS). The protein complex was then purified by Size Exclusion Chromatography (SEC), using a HiLoad 16/600 Superdex 75 pg column at 1 mL/min. The identity of the purified proteins was further confirmed in reduced and non-reduced conditions using sodium dodecyl sulfate-polyacrylamide (SDS-PAGE) gel electrophoresis.

### Expression & Purification of His6-SpyCatcher-mi3 (VLP Scaffold)

The virus-like particle (VLP) with an N-terminal 6x-His Tag, His6-SpyCatcher-mi3, was produced as previously described.^25,30,38^ The plasmid encoding the SpyCatcher-mi3 particle (Addgene 113043) was transformed into *Escherichia coli* BL21(DE3) cells. A starter culture of 50 mL in LB Broth supplemented with of 50 μg/mL kanamycin was done overnight (16 hours). The starter culture was then diluted 1:100 into 1L of LB supplemented with 50 μg/mL kanamycin and 0.8% (w/v) glucose; this was cultured at 37°C at 200 rpm until an OD600 of 0.8 was reached. Protein expression was induced with isopropyl β-D-1-thiogalactopyranoside (IPTG) (420 μM) and incubated at 18°C at 200 rpm for an additional 16-18 hours. The culture was centrifuged at 6000 rpm for 20 minutes at 4°C. The resulting pellet was resuspended in 30 mL buffer containing 250 mM Tris-HCl, 150 mM NaCl, 50 mM imidazole, 0.02% sodium azide (NaN_3_), supplemented with 0.1 mg/mL lysozyme, 2 mM phenylmethanesulfonyl fluoride (PMSF), and 1000X benzonase nuclease. Cell suspension was incubated at room temperature for 1 hour on a platform shaker before being sonicated on ice five times of 30-second intervals at 40% amplitude (30 seconds on followed by 30 seconds off). Cell lysate was then clarified at 21,000 × g for 30 minutes at 4°C. The supernatant was filtered through 0.45 μm then 0.22 μm syringe filters before being isolated by Ni-NTA chromatography using a pre-packed HisPrep FF 16/10 column. For this IMAC purification, the binding buffer was 20 mM Tris-HCl pH 8.0, 150 mM NaCl, 50 mM imidazole, and 0.02% NaN_3_; SpyCatcher-mi3 particle was eluted with the elution buffer composed of 20 mM Tris-HCl pH 8.0, 150 mM NaCl, 2M imidazole, and 0.02% NaN_3_. Eluted particles were concentrated using a 30 kDa concentrator and further purified by SEC using a HiLoad® 16/600 Superdex® 200 column pre-equilibrated with 25 mM Tris-HCl pH 8.0, 150 mM NaCl, and 0.02% NaN_3_. SpyCatcher-mi3 particles were stored at 4°C and used for conjugations for up to 1 month after filtering with a 0.2 μm filter or centrifuging at 14,000 rpm for 15 minutes. The concentration of SpyCatcher-mi3 particles was measured by Pierce™ BCA Protein Assay (Thermo Scientific, 23225).

### Conjugation of SpyTag Open HLA to SpyCatcher-mi3

Prior to the conjugation, the proteins were centrifuged at 15,000 rpm for 10 minutes to remove any potential aggregates. VLP-SpyCatcher-mi3 was quantified using Pierce™ BCA Protein Assay. For a 500 μL-reaction or 1000 μL-reaction, 50 nM of purified VLP-SpyCatcher-mi3 (molarity of 60-mer particle) was incubated with 9 μM SpyTag Open HLA-A*02:01/NY-ESO-1^C9V^, SpyTag Open HLA-A*02:01/NY-ESO-1^W5A^, or SpyTag Open HLA-A*02:01/gTAX in 25 mM Tris-HCl pH 8.0, 150 mM NaCl, 0.02% NaN_3_ at room temperature for 16-18 hours. The VLP-SpyCatcher particle:SpyTag molar ratio explored was 1:180 as optimized. Coupling efficiency was visualized with SDS-PAGE with Coomassie staining. Conjugated VLP-Open HLA complex was separated from unconjugated HLA and monomers by washing the reactions three times with 25 mM Tris-HCl pH 8.0, 150 mM NaCl, 0.02% NaN_3_ using a 100-kDa MWCO Amicon® Ultra Centrifugal Filter spin column. The concentration of VLP-Open HLA complex was measured by Pierce™ BCA Protein Assay.

### Dynamic Light Scattering (DLS)

For each sample, VLP-Open HLA protein complex in buffer (25 mM Tris-HCl pH 8.0, 150 mM NaCl, and 0.02% NaN_3_) were centrifuged at 15,000 rpm for 10 minutes to remove any potential aggregates right before DLS experiment. In a cuvette, 60 μL of each sample was loaded; the total protein concentration should be between 0.2-0.9 mg/mL. Each sample was measured at 20°C on a Nanobrook 90Plus Submicron Particle Sizer (Brookhaven Instruments; Nashua, NH, USA) for 2-4 runs. The intensity distribution was plotted and the particle size/radius average along with its standard deviation were calculated in Excel.

### Mass Photometry

Prior to characterization with mass photometry, the VLP particles (either conjugated or not conjugated with Open HLA) were centrifuged at 15,000 rpm for 15 minutes at 4°C. The stock particles were stored in a buffer of 25 mM Tris-HCl pH 8.0, 150 mM NaCl, and 0.02% NaN_3_, and they could be further diluted in 1X PBS to the final concentrations right before data acquisition. The mass photometry instrument was tested with sterile 1X PBS buffer and calibrated with a mass standard of β-amylase (112, 224 kDa) and thyroglobulin (670 kDa). The particles of concentration between 200-1000 μg/mL were examined by localizing 1-2 μL/sample on the mass photometry grid. The data were collected on a TwoMP mass photometer (Refeyn; Oxford, United Kingdom) and analyzed with the Refeyn Discover MP analysis software in histogram mode and mass plot using default settings. For each sample, an automated Gaussian curve was fitted to the particle distribution to estimate the molecular counts, median masses, and their standard deviation (sigma values). Mass photometry data for all particles are shown in Supporting Information, **Figure S4**.

### Cryo EM

VLP-SpyCatcher-mi3 was conjugated to SpyTag Open HLA-A*02:01/NY-ESO-1^C9V^ using the method in the previous section. The particle VLP-Open A*02:01/NY-ESO-1^C9V^ at a total protein concentration of 0.919 mg/mL was used for grid preparation. The sample was prepared on Quantifoil Cu 300 2/2 (Quantifoil) grids and was frozen in liquid ethane using the Vitroblot Mark IV (Thermo Fisher) at 4°C and 100% humidity. The micrographs were collected on a Titan Krios G3i 300 kV cryo electron microscope (Thermo Fisher Scientific) with a K3 Summit camera (Gatan).

### Fluorescence Polarization (FP)

VLP-Open HLA-A*02:01/gTAX conjugation was conducted in the same way as described in the previous section. Open A*02:01 was refolded with the placeholder peptide, gTAX, was exchanged and loaded with labeled peptide _TAMRA_TAX9 (TAMRA-KLFGYPVYV). Prior to the assays, the VLPs and all components of the assays were centrifuged for 15,000 rpm; VLP concentration was determined via Pierce™ BCA Protein Assay. In a 96-well plate, for each 100 μL reaction, VLP-Open HLA-A*02:01/gTAX of different concentrations (2.5-100 nM) were incubated with 40 nM _TAMRA_TAX9 in FP buffer (150 mM NaCl, 20 mM sodium phosphate, 0.05% Tween-20, pH 7.4). The concentration of the TAMRA-labeled peptide was previously optimized so that the polarization baseline was between 0 and 50 mP. The kinetic association of fluorescently labeled peptides loaded on MHC-I was monitored via FP for 4-6 hours. The fluorescence of the TAMRA-labeled peptides was monitored at the excitation and emission wavelengths of 531 and 595 nm on a SpectraMax iD5 plate reader. All experiments were performed in triplicates at room temperature. Raw parallel (III) and perpendicular emission intensities (I⊥) were measured then converted to polarization (mP) values using the equation 1000*[(III-(G*I⊥))/(III+(G*I⊥))], in which the G-factor was 0.33 for TAMRA-labeled peptides. The data was fitted in GraphPad Prism 10.

### Fluorescent Labeling of VLP-Open HLA

SpyCatcher-mi3 (VLP) was centrifuged at 16,000 rpm for 15 minutes at 4°C to remove any potential aggregate. The particle was previously quantified using Pierce™ BCA Protein Assay. The particle was dialyzed into 50 mM borate buffer pH 8.5, to ensure that there is no sodium azide left as sodium azide can interfere with the labeling reaction. After being dialyzed overnight, the particles were concentrated using a 10kDa spin column preequilibrated with 50 mM borate buffer pH 8.5. The particle concentration was then re-quantified using Pierce™ BCA Protein Assay to ensure accurate calculation for the labeling reaction, and the VLP particles were diluted to the concentration of 2 mg/mL. NHS-fluoresceine (Thermo Scientific, 46410; molecular weight 473.4 g/mol) was equilibrated to room temperature about 20 minutes before the labeling reaction to avoid moisture. NHS-fluoresceine is reconstitute in DMSO (1 mg/100 μL). The amount of NHS-fluoresceine was calculated such that there is 15 mmol NHS-fluoresceine per 1 mmol VLP monomer. The reaction was conducted in 50 mM borate buffer 8.5 for 2 hours on ice in an amber tube shieled from light. To remove excess dye from the reaction, the Fluorescent Dye Removal Column (Thermo Scientific, 22858) was used and the protocol from the manufacturer was followed. The purified, fluorescently labeled VLP particles were then quantified using the Pierce™ BCA Protein Assay. The samples were analyzed with MALDI-MS to confirm that the labeling reaction was successful.

### T Cell Staining with Fluoresceine Labeled VLP-Open HLA

CD8+ T cells 1G4 TCR, a human T cells engineered expressing the variants of a single TCR (1G4) specific for the cancer antigen NY-ESO-1, were used in these experiments. For details regarding the production of these cells, see the Supporting Information. The cells were cultivated in Advanced RPMI (Gibco), supplemented with 10% heat-inactivated FBS (Gibco), 1X Glutamax (Gibco), 1X Penicillin-Streptomycin (Gibco), 10 mM HEPES (Gibco), and 10 ng/mL recombinant IL-2. In a 96-well TC-treated plate, for each well, 200 μL of T cells were cultured at 2 x 10^6^ cells/mL at 37°C and 5% CO_2_ overnight. On the next day, the cells were harvested by centrifuging the plate at 300g for 5 minutes then stained with 500 µL of 2000X diluted LIVE/DEAD IR 876 Fixable Dead Cell Stains (Invitrogen) for 10 minutes at room temperature being centrifuged at 300g for 5 minutes and washed with 500 µL FACS buffer. T cells were stained on ice with 100 µL of 1 μg/mL of either tetramers (PE-Open HLA-A*02:01/NY-ESO-1^C9V^ or PE-Open HLA-A*02:01/NY-ESO-1^W5A^ tetramers) or fluorescent-labeled VLP-Open HLA VLP-Open HLA-A*02:01/NY-ESO-1^C9V^ or VLP-Open HLA-A*02:01/NY-ESO-1 ^W5A^). The samples were further stained for 20 minutes on ice with 100 µL of the diluted Brilliant Ultra Violet^TM^ 605 CD8 antibody (Biolegend, 344742) and APC anti-human TCR Vβ13.1 antibody (Biolegend, 362408). The cells were washed and centrifuged again, then fixed with 4% paraformaldehyde (PFA) for 30 minutes at room temperature. After being washed twice with FACS buffer, the cells were ready to be analyzed with a CytoFLEX LX flow cytometer. For gating strategy, see **Figure S15**.

### T Cell Activation Assay & Flow Cytometry

We conducted a T cell activation assay with VLP-Open HLA-A*02:01/NY-ESO-1^C9V^ (active construct) and VLP-Open HLA-A*02:01/NY-ESO-1 ^W5A^ (inactive construct as a negative control). CD8+ T cells 1G4 TCR were used in these experiments. The cells were cultivated in Advanced RPMI (Gibco), supplemented with 10% heat-inactivated FBS (Gibco), 1X Glutamax (Gibco), 1X Penicillin-Streptomycin (Gibco), 10 mM HEPES (Gibco), and 10 ng/mL recombinant IL-2.

On Day 0 of the experiment, T cells were cultured at 1 x 10^6^ cells/mL at 37°C and 5% CO_2_ overnight. On Day 1, the cells were plated at 100,000 cells/well in a sterile U-bottomed 96-well plate; the cells could be stained with CellTrace Violet proliferation dye (Thermo) prior to stimulation according to the manufacturer’s protocol. The cells were stimulated with different concentrations of the VLP particles (VLP-Open HLA-A*02:01/NY-ESO-1^C9V^ or VLP-Open HLA-A*02:01/NY-ESO-1 ^W5A^) or at a 1:1 ratio of Dynabeads™ Human T-Activator CD3/CD28 for T Cell Expansion and Activation (Gibco) for 72 hours (3 days). A negative control with media only (no VLP and no beads) was also included. For the conditions with VLPs and the negative control, the media was also supplemented with 4 μg/mL CD28. Each condition was done in duplicates. On Day 3, harvest the cells by centrifuging the plate at 300g for 5 minutes. The cell supernatant could be saved for later analysis with ELISA. For flow cytometry analysis, the cells were stained with 500 µL of 2000X diluted LIVE/DEAD IR 876 Fixable Dead Cell Stains (Invitrogen) for 10 minutes at room temperature being centrifuged at 300g for 5 minutes and washed with 500 µL FACS buffer. The cells were centrifuged again at 300g for 5 minutes then stained with 100 µL of an antibody cocktail (Brilliant Ultra Violet™ 661 or 650 Dye* for CD69; FITC mouse anti-human for CD8, PE/Cyanine7 anti-human for CD137 or 4-1BB) in 4°C fridge for 30 mins. The cells were washed and centrifuged again, then fixed with 4% PFA for 30 minutes at room temperature. After being washed twice with FACS buffer, the cells were ready to be analyzed with a CytoFLEX LX flow cytometer.Other antibodies used for surface staining were anti-PD-1 PE-Dazzle-594 (Clone: EH12.2H7) and anti-CD8 BV605 (Clone: SK1).

For intracellular staining for Tbet and Eomes, cells were fixed with Fix and PERM Medium A (Thermo Fisher) for 20 minutes, washed twice and then permeabilized with Fix and PERM Medium B(Thermo Fisher) containing Anti-T-bet PE-Cy5 (Clone: 4B10) and anti-Eomes PE-Cy7 (Clone: WD1928). For intracellular cytokine staining cells were stimulated with 1x Cell Activation Cocktail (Biolegend; containing PMA, ionomycin and brefeldin A) for 3 hours before fixation and permeabilization as above and overnight incubation with anti-IL-2 BV650 (Clone: MQ1-17H12) and anti-IFN-γ BV510 (Clone: B27). All permeablized cells were washed twice with BD Perm Wash buffer (BD Biosciences) twice before one additional wash in FACS buffer and acquired on a Aurora flow cytometer (Cytek).

For gating strategy, we identified singlets from FSC-A vs. FSC H, then identified live cells from FSC-A vs. Live-Dead IR 876-A. From this subpopulation of cells, we identified CD8+ by gating FSC-A vs. CD8+. The CD8+ cell subpopulation was gated against 4-1BB to identify the population expressing 4-1BB, a marker of T cell activation (Supplemental Information, **Figure S16**).

### ELISA

The cell supernatant from T cell activation assays were stored in -20°C freezer until analysis by ELISA. The samples were diluted 2X-3X for ELISA analysis with DuoSet ELISA: Human IFN-γ (R&D-Biotechne; DY285B) and DuoSet Ancillary Reagent Kit (R&D – Biotechne; DY008) according to the manufacturer’s protocol. In brief, a flat-bottom polystyrene microplate was coated with 2.00 μg/mL capture antibody overnight before being washed and blocked with the Reagent Diluent containing 1% BSA. In addition to the samples, a standard curve with recombinant human IFN-γ was prepared in Reagent Diluent was prepared, and they were added to the plate with an incubation time of 2 hours at room temperature. The plate was washed with the Wash Buffer before adding Detection Antibody (biotinylated mouse anti-human IFN-γ) at 200 ng/mL for another 2-hour incubation at room temperature. The plate was washed and 1X Streptavidin-HRP B was added to the plate for 20 minutes incubation at room temperature. The plate was washed again before TMB ELISA Substrate was added to each well for a 20-minute incubation at room temperature. A stop solution (2N sulfuric acid) was added to each well to stop the reaction. The OD at 450 nm (correction at 540 nm or 570 nm) was collected using a SpectraMax iD5 plate reader.

## Supporting Information

The supporting information contains additional experimental details, including information regarding peptide & protein sequences, recombinant protein purification, differential scanning fluorimetry (DSF), conjugation optimization, dynamic light scattering (DLS), mass photometry, fluorescence polarization (FP), mass spectrometry data, T cell activation study, and ELISA assays.

## Author Contribution

Conceptualization: NGS

Methodology: HATP, MCY, DH

Investigation: HATP, DH, SMS

Visualization: HATP, DH

Funding acquisition: NGS, DM

Supervision: NGS

Writing – original draft: HATP, NGS

Writing – review & editing: HATP, DH, NGS

## Notes

NGS and DH are listed as co-inventors in provisional patent application related to the use of VLP nanoparticles to display class I and class II HLA molecules, for various applications.

## Acknowledgements

This project was funded in part by the National Institute of Allergy and Infectious Diseases, National Institutes of Health, Department of Health and Human Services (NIH R01AI143997 to N.G.S.). Further funding was provided by NIH NIGMS R35GM125034 (to N.G.S.), and The Children’s Hospital of Philadelphia Cell and Gene Therapy Collaborative. The authors are grateful to Dr. Leland Mayne for assistance with mass spectrometry (LC-MS), Ryan Kubanoff for assistance with Bruker Ultraflex III matrix-assisted laser desorption/ionization time-of-flight mass spectrometer (MALDI-TOF/TOF MS) in the University of Pennsylvania’s Chemistry Department (instrument grant NIH S10-OD030460), Dr. Ronen Marmorstein for mass photometry, Dr. Leena Mallik and the University of Pennsylvania’s Beckman Center for Cryo-Electron Microscopy for assistance with our cryo-EM data collection and analysis.

## 1. Peptides & Ligands

The peptides for this study were purchased from GenScript (Piscataway, NJ, USA) at > 90% purity. TAMRA-TAX9, the TAX9 peptide fluorescently labeled with 5-carboxytetramethylrhodamine (TAMRA), was purchased from Biopeptek Pharmaceuticals LLC (Malvern, PA, USA). Peptides were solubilized in Milli-Q water and centrifuged at 14,000 rpm for 15 minutes to remove any insoluble peptide. The peptide concentration was measured and quantified using the absorbance and extinction coefficient at 205 nm. The peptide information, with peptide sequences given in standard single-letter codes, is shown in **Table S1**. For the TCGA mass spectrometry peptide study, the peptides were selected based on our previous study (Sun *et al.,* 2023)^1^ and the peptides were purchased from Mimotopes (Australia); peptide information for the TCGA peptide is shown in **Table S4**.

**TABLE S1.**
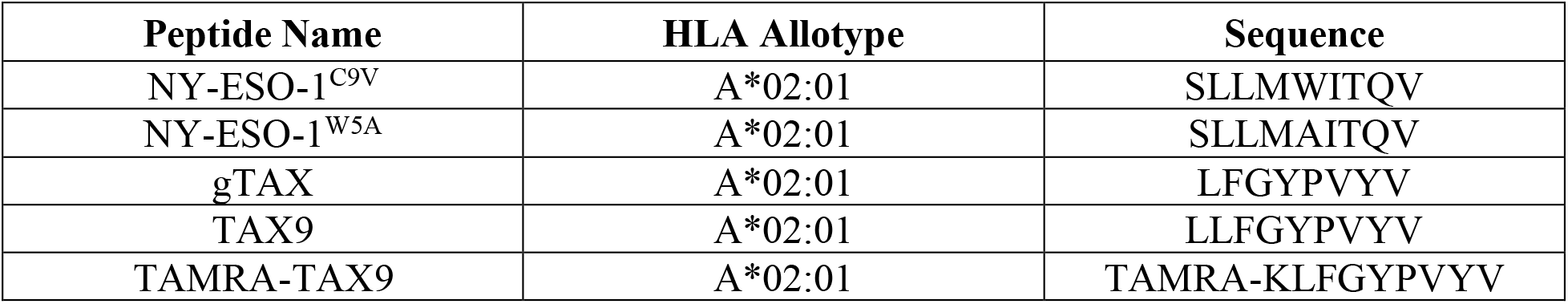
Sequence information of the peptides used in this study.

| Peptide Name | HLA Allotype | Sequence |
| --- | --- | --- |
| NY-ESO-1<sup>C9V</sup> | A*02:01 | SLLMWITQV |
| NY-ESO-1<sup>W5A</sup> | A*02:01 | SLLMAITQV |
| gTAX | A*02:01 | LFGYPVYV |
| TAX9 | A*02:01 | LLFGYPVYV |
| TAMRA-TAX9 | A*02:01 | TAMRA-KLFGYPVYV |

## 2. Recombinant Protein, Refolding, and Purification Plasmid Information & Protein Sequences

The plasmids were purchased from either GenScript (Piscataway, NJ, USA) or Addgene (Watertown, MA, USA). The protein sequences for the proteins used in this study along with their plasmid sources is included in **Table S2** below.

**TABLE S2.**
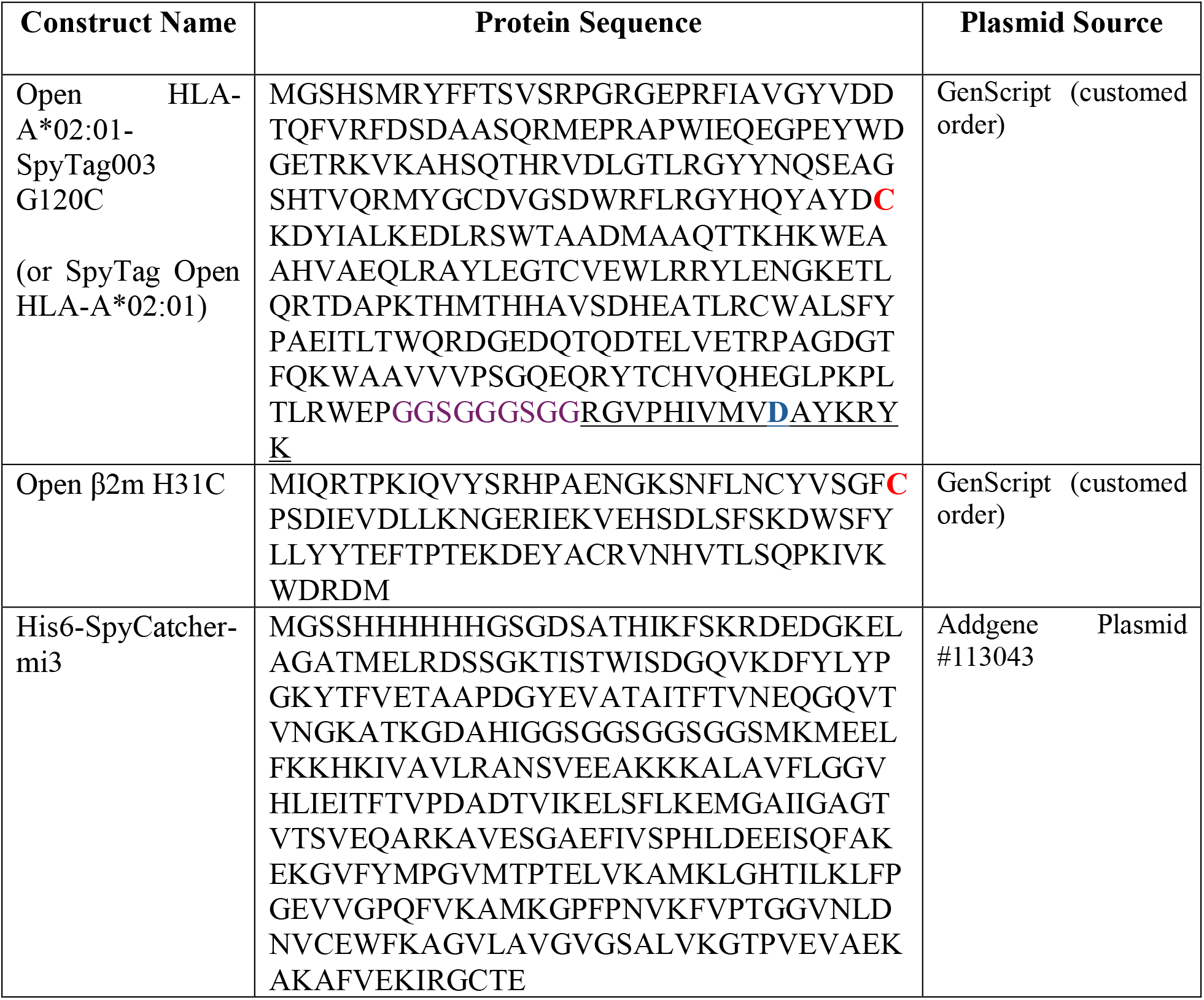
Protein Sequences.

### Expression & Purification of SpyTag Open HLA-A*02:01

The heavy chain of Open A*02:01 (G120C) ectodomain was C-terminally tagged with SpyTag003 (RGVPHIVMVDAYKRYK). This heavy chain was refolded with β2m (H31C) in the presence of excess peptide NY-ESO-1^C9V^ (SLLMWITQV), NY-ESO-1^W5A^ (SLLMAITQV; used negative control), or gTAX (LFGYPVYV; placeholder peptide). A Gly-rich linker (GGSGGGSGG) was included between the isopeptide fusion tag (SpyTag003) and the α3 domain of the heavy chain, to prevent interference between the tag function and the heavy chain and other undesired interactions. Plasmids encoding the luminal domain of the C-terminally linked SpyTag003 Open A*02:01 (G120C) and open human β2m (H31C) were transformed into *Escherichia coli* BL21(DE3) cells using the pET22b(+). The specific protein sequences are described in the Supporting Information. The cells were grown and harvested in the Luria-Broth (LB) medium, and inclusion bodies were pelleted and purified as previously described.^1^ The ternary pMHC-I complexes were generated by an *in vitro* refolding, by slowly diluting a 100 mg of Open HLA-A*02:01-SpyTag003 and open β2m at a 1:3 molar ratio over 4 hours in 500 mL of refolding buffer (0.4 M L-Arginine HCl, 100 mM Tris, 2 mM ethylenediaminetetraacetic acid (EDTA), 5 mM reduced L-glutathione, 0.5 mM oxidized L-glutathione, pH 8.0) supplemented with 5 mg of the peptide. The refolding reaction proceeded for 4 days, and the proteins were then dialyzed into a buffer containing 25 mM Tris, 150 mM NaCl, pH 8.0 (TBS). The protein complex was then purified by Size Exclusion Chromatography (SEC), using a HiLoad 16/600 Superdex 75 pg column at 1 mL/min. The identity of the purified proteins was further confirmed in reduced and non-reduced conditions using sodium dodecyl sulfate-polyacrylamide (SDS-PAGE) gel electrophoresis.

### Expression & Purification of His6-SpyCatcher-mi3 (SpyCatcher-VLP Scaffold)

The virus-like particle (VLP) with an N-terminal 6x-His Tag, His6-SpyCatcher-mi3, was produced as previously described.^2–4^ The plasmid encoding the SpyCatcher-mi3 particle (Addgene 113043) was transformed into *Escherichia coli* BL21(DE3) cells. A starter culture of 50 mL in LB Broth supplemented with of 50 μg/mL kanamycin was done overnight (16 hours). The starter culture was then diluted 1:100 into 1L of LB supplemented with 50 μg/mL kanamycin and 0.8% (w/v) glucose; this was cultured at 37°C at 200 rpm until an OD600 of 0.8 was reached. Protein expression was induced with isopropyl β-D-1-thiogalactopyranoside (IPTG; 420 μM) and incubated at 18°C at 200 rpm for an additional 16-18 hours. The culture was centrifuged at 6000 rpm for 20 minutes at 4°C. The resulting pellet was resuspended in 30 mL buffer containing 250 mM Tris-HCl, 150 mM NaCl, 50 mM imidazole, 0.02% sodium azide (NaN_3_), supplemented with 0.1 mg/mL lysozyme, 2 mM phenylmethanesulfonyl fluoride (PMSF), and 1000X benzonase nuclease. Cell suspension was incubated at room temperature for 1 hour on a platform shaker before being sonicated on ice five times of 30-second intervals at 40% amplitude (30 seconds on followed by 30 seconds off). Cell lysate was then clarified at 21,000 × g for 30 minutes at 4°C. The supernatant was filtered through 0.45 μm then 0.22 μm syringe filters before being isolated by Ni-NTA chromatography using a pre-packed HisPrep FF 16/10 column. For this immobilized metal affinity chromatography (IMAC) purification, the binding buffer was 20 mM Tris-HCl pH 8.0, 150 mM NaCl, 50 mM imidazole, and 0.02% NaN_3_; the elution buffer was composed of 20 mM Tris-HCl pH 8.0, 150 mM NaCl, 2M imidazole, and 0.02% NaN_3_ (**Figures S1A-1B**). Eluted particles were concentrated using a 30 kDa concentrator and further purified by SEC using a HiLoad® 16/600 Superdex® 200 column pre-equilibrated with 25 mM Tris-HCl pH 8.0, 150 mM NaCl, and 0.02% NaN_3_ (**Figures S1C-1D)**. SpyCatcher-mi3 particles were stored at 4°C and used for conjugations for up to 1 month after filtering with a 0.2 μm filter or centrifuging at 14,000-15,000 rpm for 15 minutes.

**Figure S1.**
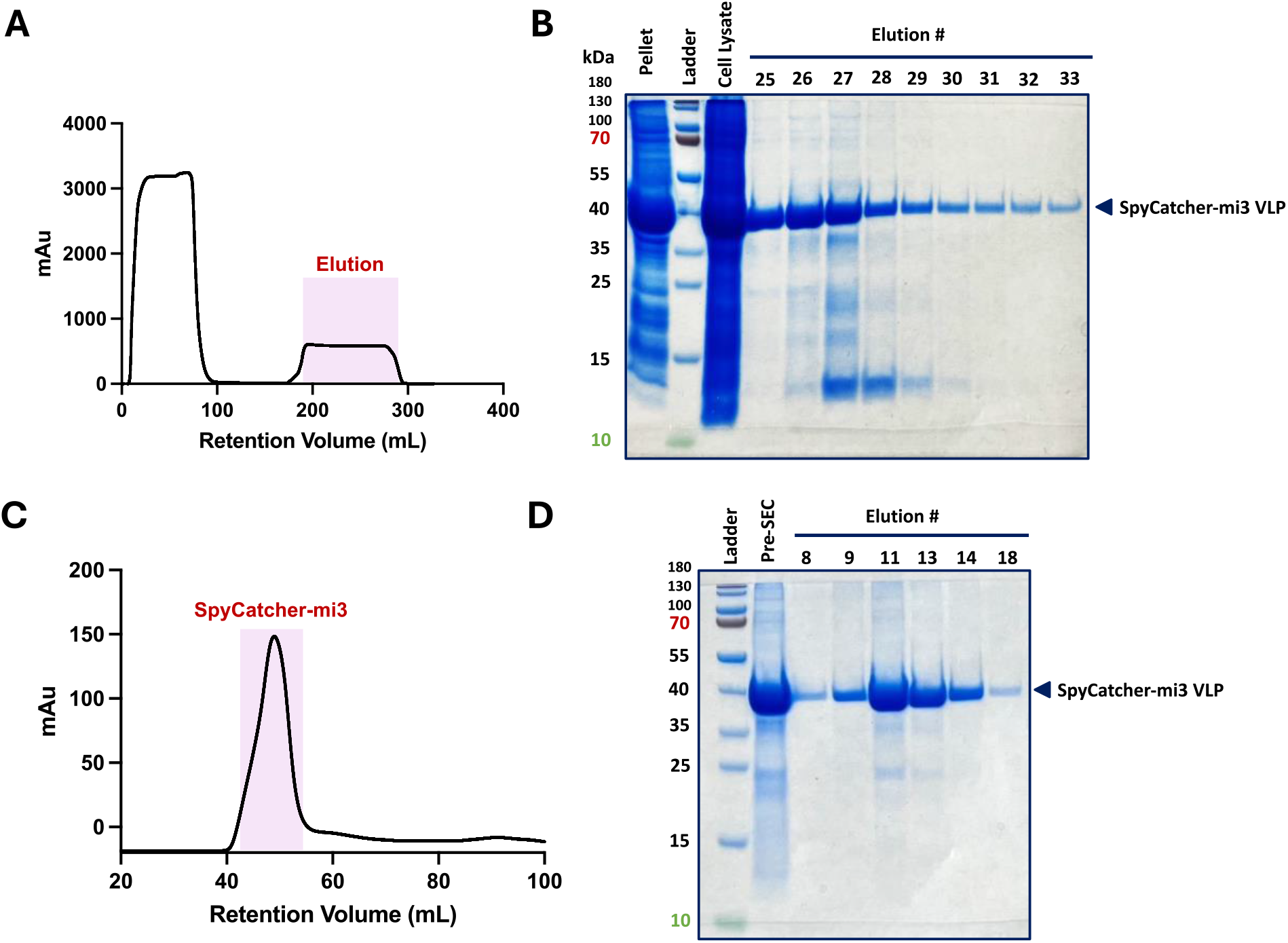
Expression and Purification of His6-SpyCatcher-mi3 (SpyCatcher-VLP Scaffold). **(A)** IMAC trace of His6-SpyCatcher-mi3. **(B)** SDS-PAGE of fractions corresponded to the highlighted elution peak from the IMAC trace. **(C)** SEC profile of His6-SpyCatcher-mi3. **(D)** SDS-PAGE analysis of fractions corresponded to the highlighted elution peak from the SEC trace.

### VLP Particle Concentration Quantification

Using the manufacturer’s established protocol, the concentration of SpyCatcher-mi3 particles, either unconjugated or conjugated with SpyTag HLA proteins, was measured by Pierce™ BCA Protein Assay (Thermo Scientific, Catalog #23225).

## 3. Differential Scanning Fluorimetry (DSF)

Differential Scanning Fluorimetry (DSF) was conducted to evaluate the thermal stability of the following pMHC-I complexes: SpyTag Open HLA-A*02:01 refolded with either NY-ESO-1^C9V^ (SLLMWITQV), NY-ESO-1^W5A^ (SLLMAITQV; negative control), or gTAX (LFGYPVYV; placeholder peptide). To a final volume of 20 μL, the pMHC-I complex at 7 μM final concentration was mixed thoroughly with 10X SYPRO^TM^ Orange Protein Stain in a buffer comprised of 50 mM NaCl, 20 mM sodium phosphate, pH 7.2. Samples were prepared in triplicates then loaded into MicroAmp™ Optical 384-Well Reaction Plate (Applied Biosystems™, Thermo Fisher Scientific, Catalog #4309849). The plate was sealed with MicroAmp™ Optical Adhesive Film (Applied Biosystems™, Thermo Fisher Scientific, Catalog #4360954) and was spun at 1000 rpm for 1 minute prior to being loaded into the instrument. The samples were read at an excitation wavelength of 470 nm and an emission wavelength of 569 nm, using a QuantStudio^TM^ 5 Real-Time PCR system. The temperature was incrementally increased from 25°C to 95°C at a constant rate of 1°C per minute. The raw fluorescence intensity was normalized against the maximum intensity. Data analysis was performed with GraphPad Prism 10, where the thermal melting temperature was obtained via a nonlinear fit using the Boltzmann sigmoidal equation.

**Figure S2.**
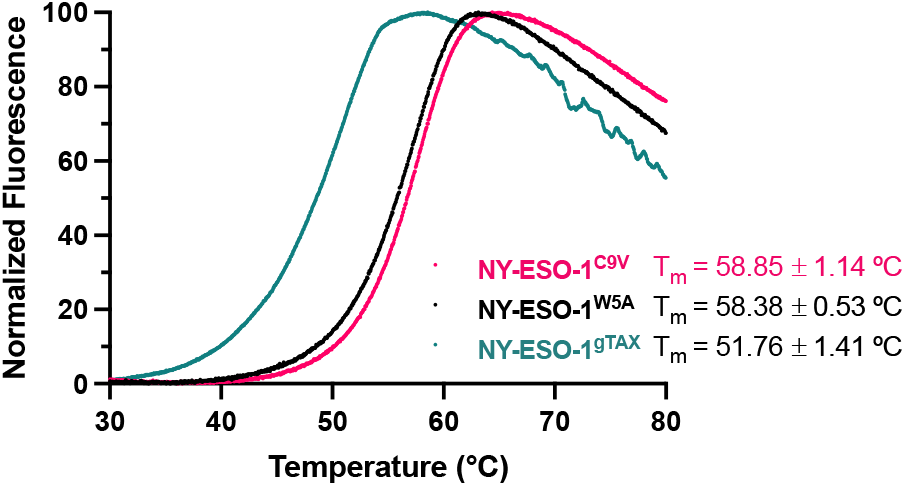
DSF thermal stability curves of SpyTag HLA complexes. Normalized fluorescence intensity is shown for SpyTag Open HLA-A*02:01 refolded with either NY-ESO-1^C9V^ (pink curve; T_m_ = 58.85°C), NY-ESO-1^W5A^ (black curve; T_m_ = 58.38°C), or gTAX (green curve; T_m_ = 51.76°C). The average (mean) of three technical replicates is plotted.

## 4. SpyCatcher-mi3 and SpyTag Open HLA Conjugation Optimization

To saturate the binding sites (60 binding sites on VLP) and improve the efficiency of VLP conjugation to open HLA molecules, micro-reactions of 20 μL/reaction were done to optimize the conjugation reaction. For each reaction, 50 nM of purified SpyCatcher-mi3 (SpyCatcher-VLP) was incubated with different molar ratio of SpyTag open HLA-A*02:01/NY-ESO-1^C9V^ or open HLA-A*02:01/NY-ESO-1^W5A^ (**Table S3**) in 25 mM Tris-HCl pH 8.0, 150 mM NaCl, 0.02% NaN_3_ at room temperature for 16-18 hours. The SpyCatcher-VLP:SpyTag HLA molar ratio explored was 1:60, 1:120, and 1:180. Coupling efficiency was visualized with SDS-PAGE with Coomassie staining (**Figure S3**). The concentration of SpyCatcher-VLP pre-conjugation and post-conjugation was measured by Pierce™ BCA Protein Assay (Thermo Scientific, Catalog #23225). Our results suggested that a ratio of 1:180, with final concentrations such as 50 nM SpyCatcher-VLP and 9 μM SpyTag Open HLA resulted in the best conjugation reaction.

**Table S3.** Equivalent (VLP:HLA molar ratio) and concentration for the conjugation reactions. Each SpyCatcher-VLP particle can accommodate up to 60 conjugation sites.

|  | *Equivalent (VLP:HLA Molar Ratio) & Concentration* |  |  |
| --- | --- | --- | --- |
|  | **1:60** | **1:120** | **1:180** |
| SpyCatcher-VLP | 50 nM<br>(3 μM monomer) | 50 nM<br>(3 μM monomer) | 50 nM<br>(3 μM monomer) |
| SpyTag Open HLA | 3 μM | 6 μM | 9 μM |

**Figure S3.**
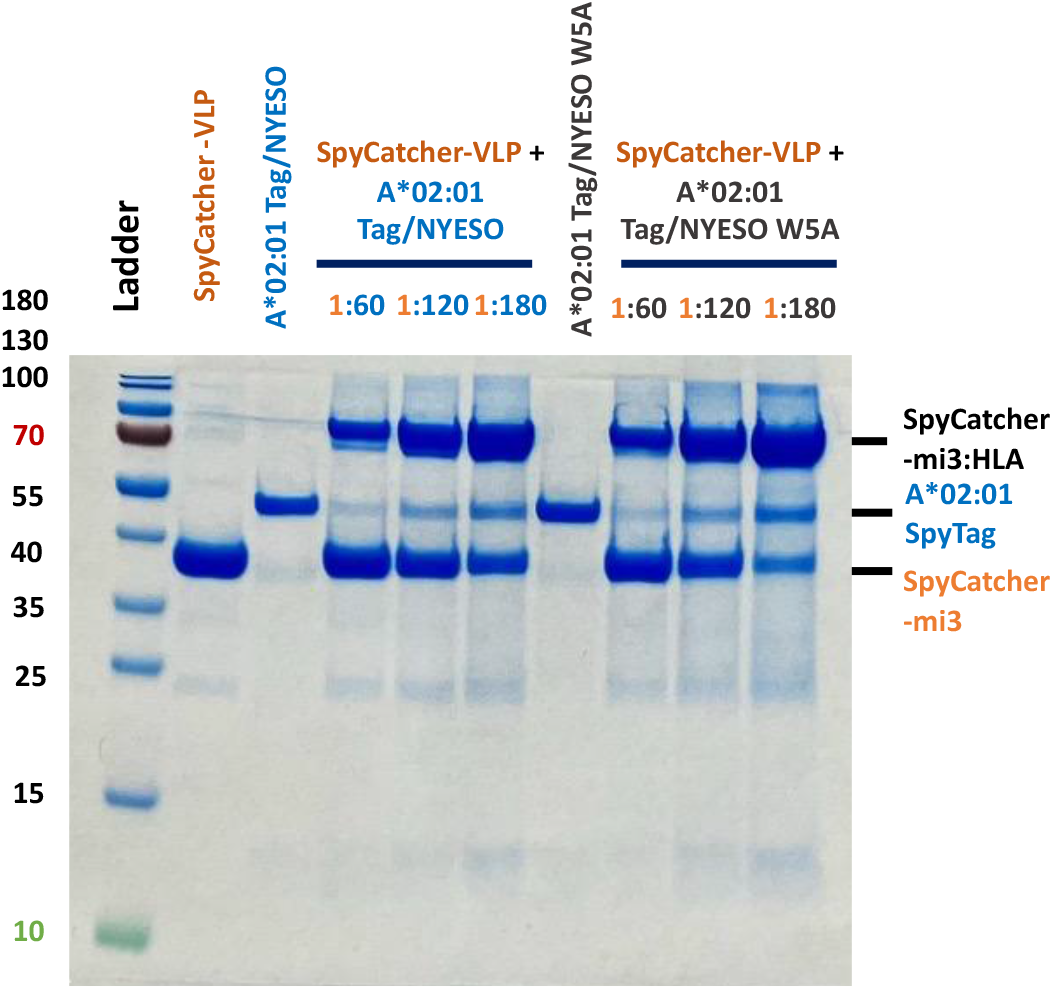
SDS-PAGE of the micro-conjugation reactions of different molar ratios between SpyCatcher-mi3 (SpyCatcher-VLP) with SpyTag open HLA-A*02:01/NY-ESO-1^C9V^ or SpyTag Open HLA-A*02:01/NY-ESO-1^W5A^. The gel showed pre- and post-conjugation products. In SDS-PAGE, the 60-mer complex broke apart and was observed as a monomer band.

## 6. Mass Photometry (MP)

Prior to characterization with mass photometry, the VLP particles (either conjugated or not conjugated with Open HLA) were centrifuged at 15,000 rpm for 15 minutes at 4°C. The stock particles were stored in a buffer of 25 mM Tris-HCl pH 8.0, 150 mM NaCl, and 0.02% NaN_3_, and they could be further diluted in 1X PBS to the final concentrations right before data acquisition. The mass photometry instrument was tested with sterile 1X PBS buffer and calibrated with a mass standard of β-amylase (112, 224 kDa) and thyroglobulin (670 kDa). The particles of concentration between 200-1000 μg/mL were examined by localizing 1-2 μL/sample on the mass photometry grid. The data were collected on a TwoMP mass photometer (Refeyn; Oxford, United Kingdom) and analyzed with the Refeyn Discover MP analysis software in histogram mode and mass plot using default settings. For each sample, an automated Gaussian curve was fitted to the particle distribution to estimate the molecular counts, median masses, and their standard deviation (sigma values) (**Figure S4**).

**Figure S4.**
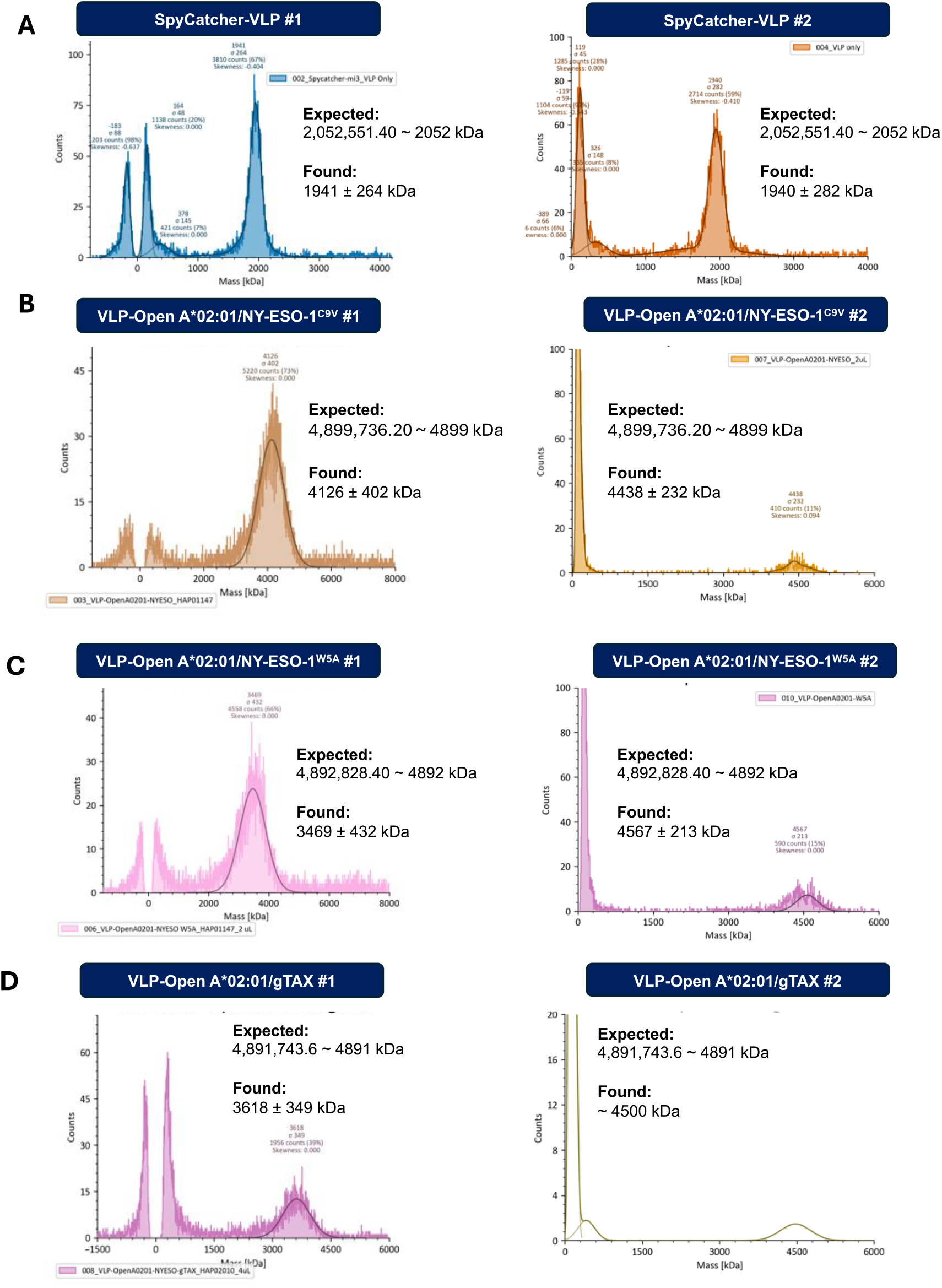
Mass photometry of unconjugated SpyCatcher-VLP and conjugated VLP-Open HLA-A*02:01 particles. Mass photometry data was obtained on two different batches of VLP particles on different experimental dates for **(A)** SpyCatcher-VLP, **(B)** VLP-Open HLA-A*02:01/NY-ESO-1^C9V^, **(C)** VLP-Open HLA-A*02:01/NY-ESO-1^W5A^, and **(D)** VLP-Open HLA-A*02:01/gTAX. The counts along with the found molecular masses and skewness values are shown.

## 7. Fluorescence Polarization (FP)

Open A*02:01 refolded with the placeholder peptide, gTAX, was conjugated to VLP to form VLP-Open HLA-A*02:01/gTAX particle. The conjugation reaction was conducted in the same way as described in the previous section. In this fluorescence polarization (FP) experiment, VLP-Open A*02:01 refolded with the placeholder peptide, gTAX, was exchanged and loaded with labeled peptide _TAMRA_TAX9 (TAMRA-KLFGYPVYV). Prior to the assays, the VLPs and all components of the assays were centrifuged for 15,000 rpm; VLP concentration was determined via Pierce™ BCA Protein Assay. In a 96-well plate, for each 100 μL reaction, VLP-Open HLA-A*02:01/gTAX of different concentrations (2.5-100 nM) were incubated with 40 nM _TAMRA_TAX9 in FP buffer (150 mM NaCl, 20 mM sodium phosphate, 0.05% Tween-20, pH 7.4). The concentration of the TAMRA-labeled peptide was previously optimized so that the polarization baseline was between 0 and 50 mP. The kinetic association of fluorescently labeled peptides loaded on MHC-I was monitored via FP for 4-6 hours. The fluorescence of the TAMRA-labeled peptides was monitored at the excitation and emission wavelengths of 531 and 595 nm on a SpectraMax iD5 plate reader. The FP experiment was done in 2 biological replicates from two different batches of VLP; the data for one biological replicate is shown in the main text Figure 2B and the other one is shown in **Figure S5**. For each biological replicate, all experiments were performed in triplicates at room temperature. Raw parallel (III) and perpendicular emission intensities (I⊥) were measured then converted to polarization (mP) values using the equation 1000*[(III-(G*I⊥))/(III+(G*I⊥))], in which the G-factor was 0.33 for TAMRA-labeled peptides. The data was fitted in GraphPad Prism 10.

**Figure S5.**
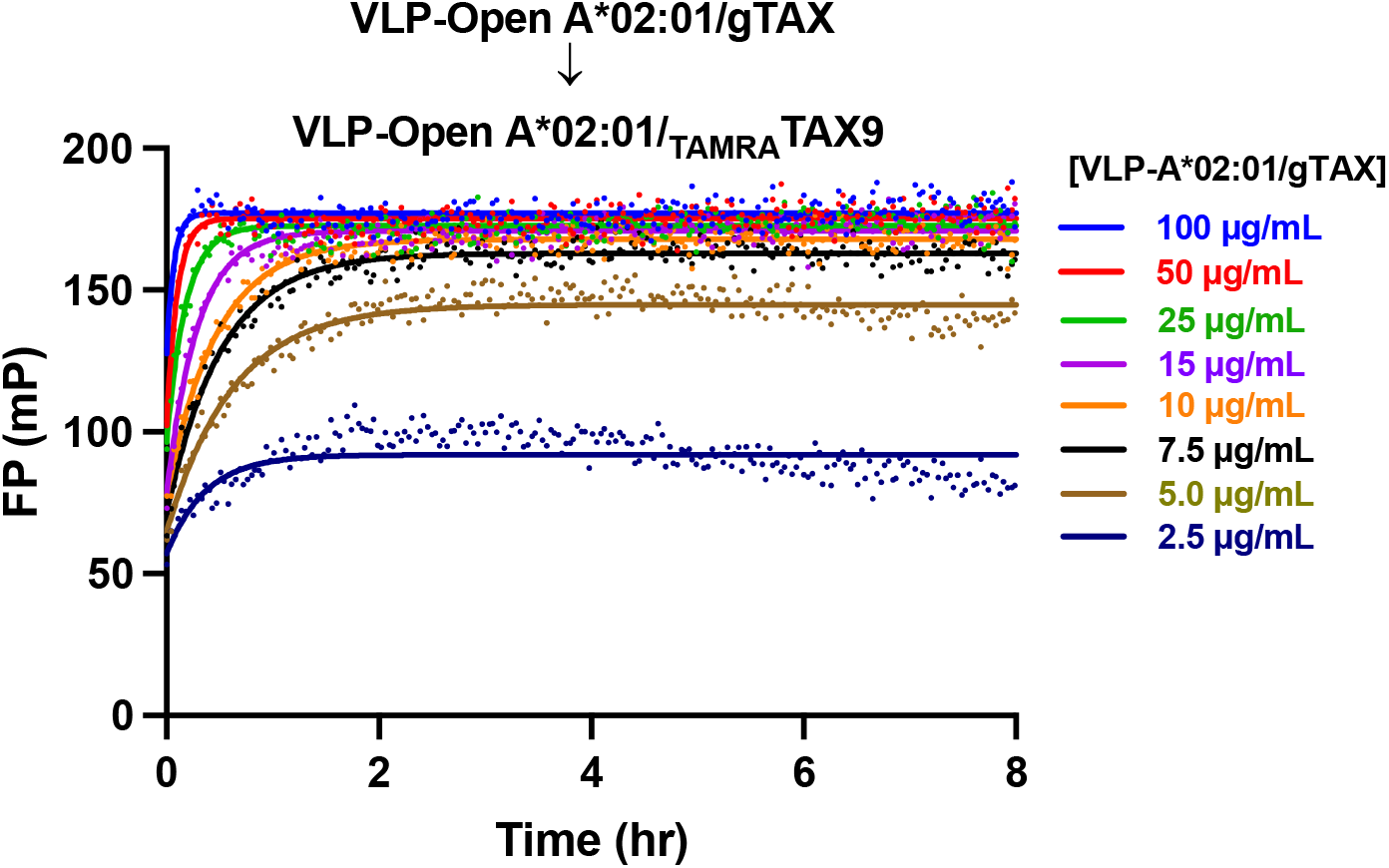
Loading and Binding of _TAMRA_TAX9 by VLP-Open A*02:01/gTAX monitored at room temperature. Peptide exchange was measured by fluorescence polarization (mP) of 40 nM _TAMRA_TAX9 as a function of the VLP-Open HLA-A*02:01/gTAX concentrations. Individual traces were fit to an exponential association model. The means from each condition’s triplicates were plotted.

## 8. Mass Spectrometry Study with TCGA Peptide Library

To prepare for the particles for the mass spectrometry study, in a total reaction volume of 1000 μL, 50 nM of purified His6-SpyCatcher-mi3 (SpyCatcher-VLP) was incubated with 9 μM SpyTag open HLA-A*02:01/gTAX in 25 mM Tris-HCl pH 8.0, 150 mM NaCl, 0.02% NaN_3_ at room temperature overnight (16-18 hours). Using the same protocol for conjugation and wash steps as outlined in the main text, SDS-PAGE was used to confirm conjugation, and the VLP-Open HLA-A*02:01/gTAX particles were washed three times using a 100-kDa spin column to remove excess unconjugated SpyTag open HLA molecules before being quantified by Pierce™ BCA Protein Assay (Thermo Scientific, Catalog #23225).

For the TCGA mass spectrometry peptide study, the peptides were selected based on our previous study (Sun *et al.,* 2023)^1^ and the peptides were purchased from Mimotopes (Australia); peptide information for the TCGA peptide is shown in **Table S4**. For the peptide exchange experiment, 4 nM (or 20 μg/mL) of VLP-Open HLA-A*02:01/gTAX particle was incubated without and with 2.4 μM peptide in 25 mM Tris-HCl pH 8.0, 150 mM NaCl, 0.02% NaN_3_ at room temperature for 1 hour then overnight at 4°C. Each reaction was prepared in a total volume of 500 μL; the different control and experimental conditions tested are summarized in **Table S5**.

The peptide exchanged samples were then be washed twice by centrifuging at 10,000 rpm using a 100-kDa spin column to remove any excess peptide. The resulting sample (about 50 μL) was diluted 2X for mass spectrometry experiment with liquid chromatography-mass spectroscopy (LC-MS) or with matrix-assisted laser desorption/ionization time-of-flight mass spectrometer (MALDI-TOF/TOF MS). For the LC-MS experiment, the sample was analyzed on a Waters Acquity UPLC SQD instrument, using a Waters Acquity C8 column followed by Electronspray Ionization-MS (ESI-MS) with a mass detection range of 0-2000 m/z. Analysis of LC-MS data was done with Xcalibur and UniDec. LC-MS spectra are shown in **Figures S6-S11**.

For the MALDI-TOF/TOF MS experiment, the samples were diluted 2X in 0.05% trifluoroacetic acid (TFA)/Milli-Q water, then spot on the MALDI plate with α-Cyano-4-Hydroxycinnamic Acid (CHCA) matrix. The samples were analyzed on the Bruker RapifleX (MALDI-TOF/TOF) (Billerica, MA, USA). MALDI MS spectra are shown in **Figure S12** and **Figure S13**.

**Table S4.** Information of peptides in the Cancer Genome Atlas (TCGA) epitope library.

| TCGA Peptide | Name | Peptide | Length | Predicted Mass |
| --- | --- | --- | --- | --- |
| 1 | KRAS_G12C | KLVVVGACGV | 10 | 944.18 |
| 2 | TP53_R273H | LLGRNSFEVHV | 11 | 1270.73 |
| 3 | KRAS_G12A | KLVVVGAAGV | 10 | 912.12 |
| 4 | TP53_H193R | RLIRVEGNLRV | 11 | 1324.56 |
| 5 | KRAS_G12S | KLVVVGASGV | 10 | 928.12 |
| 6 | TP53_K132N | ALNNMFCQLA | 10 | 1124.32 |
| 7 | PIK3CA_G118D | ILNREIDFA | 9 | 1090.22 |
| 8 | TP53_C141Y | KTYPVQLWV | 9 | 1133.33 |
| 9 | TP53_L194R | GLAPPQHRI | 9 | 988.14 |
| 10 | TP53_C238F | YMFNSSCMGGM | 11 | 1227.44 |
| 11 | TP53_V272M | LLGRNSFEMRV | 11 | 1321.54 |
| 12 | TP53_H193R | GLAPPQRLIRV | 11 | 1219.47 |
| 13 | ERBB2_S310F | YLSTDVGFCT | 10 | 1105.21 |
| 14 | FBXW7_R505G | VLMGHVAAVG | 10 | 953.15 |
| 15 | CTNNB1_S37F | YLDSGIHFG | 9 | 1008.08 |
| 16 | PIK3CA_G118D | KILNREIDFAI | 11 | 1331.55 |
| 17 | TP53_G245V | YMCNSSCMGV | 10 | 1094.3 |
| 18 | ATM_R337C | YSSGFCNIAV | 10 | 1060.17 |
| 19 | TP53_G266V | LLVRNSFEV | 9 | 1076.24 |
| 20 | NEF2L2_D29H | ILWRQDIHLGV | 11 | 1349.57 |
| 21 | TP53_C141Y | MFCQLAKTYPV | 11 | 1300.58 |
| 22 | TP53_G266V | LLVRNSFEVRV | 11 | 1331.55 |
| 23 | NFE2L2_D29H | IDILWRQDIHL | 11 | 1421.63 |
| 24 | KRAS_G12D | KLVVVGADGV | 10 | 956.13 |
| 25 | TP53_R248W | GMNWRPILTI | 10 | 1200.44 |
| 26 | PIK3CA_N345K | ILCATYVKV | 9 | 1009.25 |
| 27 | TP53_R273L | LLGRNSFEVLV | 11 | 1246.44 |
| 28 | NRAS_Q61L | LLDILDTAGL | 10 | 1043.2 |
| 29 | NRAS_Q61L | CLLDILDTAGL | 11 | 1146.34 |
| 30 | RAC1_P29S | FSGEYIPTV | 9 | 1012.1 |
| 31 | PIK3CA_C420R | RPLAWGNINL | 10 | 1153.32 |
| 32 | PIK3CA_G118D | ILNREIDFAI | 10 | 1203.38 |
| 33 | PIK3CA_E453K | GLKDLLNPI | 9 | 982.17 |
| 34 | CTNNB1_S37C | YLDSGIHCGA | 10 | 1035.12 |
| 35 | TP53_C141Y | FCQLAKTYPV | 10 | 1169.38 |
| 36 | NFE2L2_D29H | ILWRQDIHL | 9 | 1193.39 |
| 37 | SF3B1_K700E | GLVDEQQEV | 9 | 1016.05 |
| 38 | TP53_G266V | NLLVRNSFEV | 10 | 1190.34 |
| 39 | KRAS_G12V | KLVVVGAVGV | 10 | 940.17 |
| 40 | PIK3CA_N345K | KILCATYVKV | 10 | 1137.42 |
| 41 | ERBB2_S310F | YLSTDVGFCTL | 11 | 1218.37 |
| 42 | PIK3CA_H1047L | FMKQMNDAL | 9 | 1097.3 |
| 43 | CTNNB1_S37F | YLDSGIHFGA | 10 | 1079.15 |
| 44 | TP53_I195T | GLAPPQHLTRV | 11 | 1188.37 |
| 45 | TP53_K132N | ALNNMFCQL | 9 | 1053.25 |
| 46 | TP53_V272M | LLGRNSFEM | 9 | 1066.22 |
| 47 | PIK3CA_E453K | GLKDLLNPIGV | 11 | 1138.35 |
| 48 | TP53_C141Y | CQLAKTYPV | 9 | 1022.21 |
| 49 | MAP2K1_P124S | VLHECNSSYI | 10 | 1164.28 |
| 50 | KRAS_G13C | KLVVVGAGCV | 10 | 944.18 |

**Table S5.**
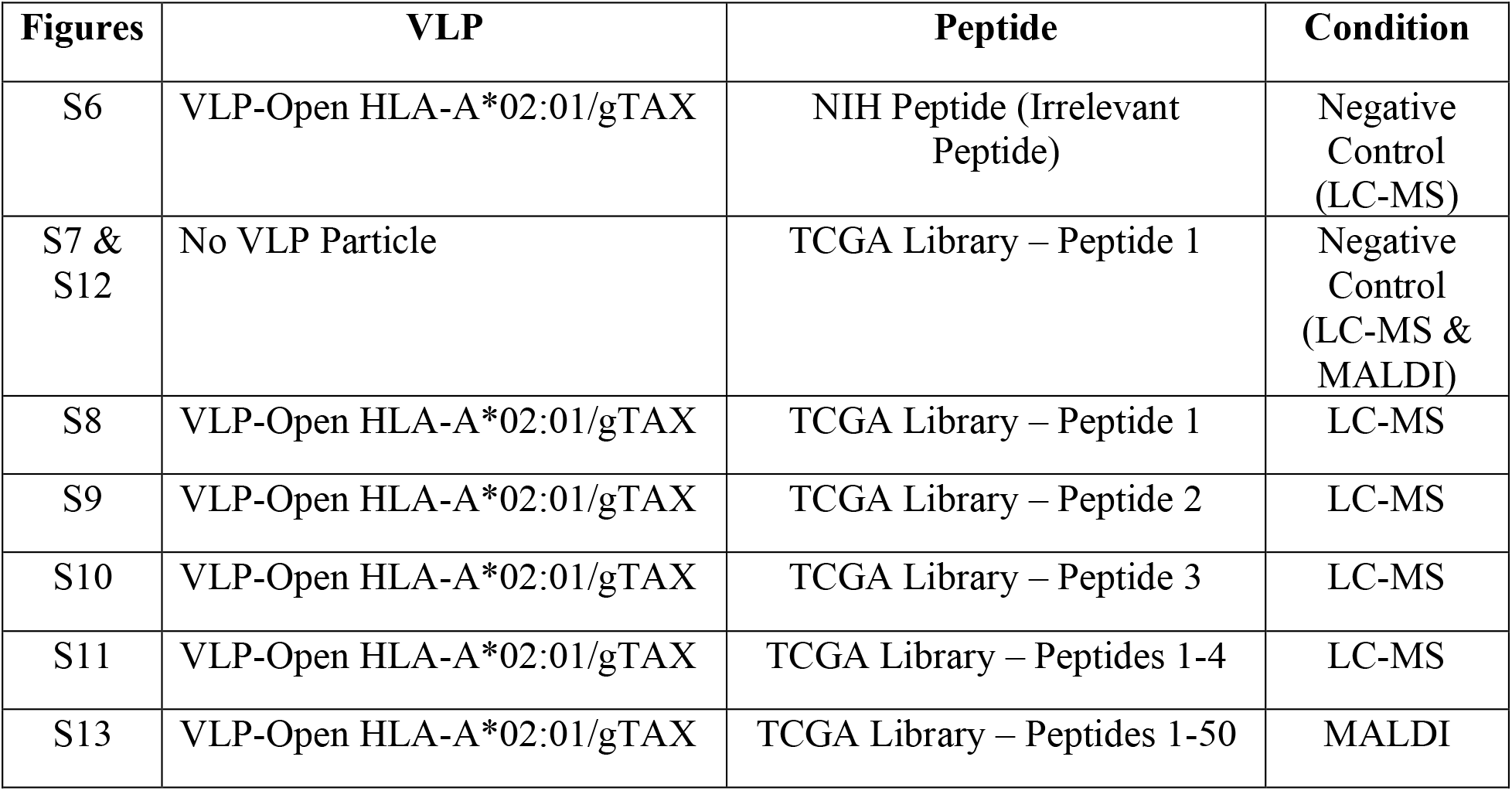
Conditions tested for mass spectrometry experiment with VLP-Open HLA particles.

**Figure S6.**
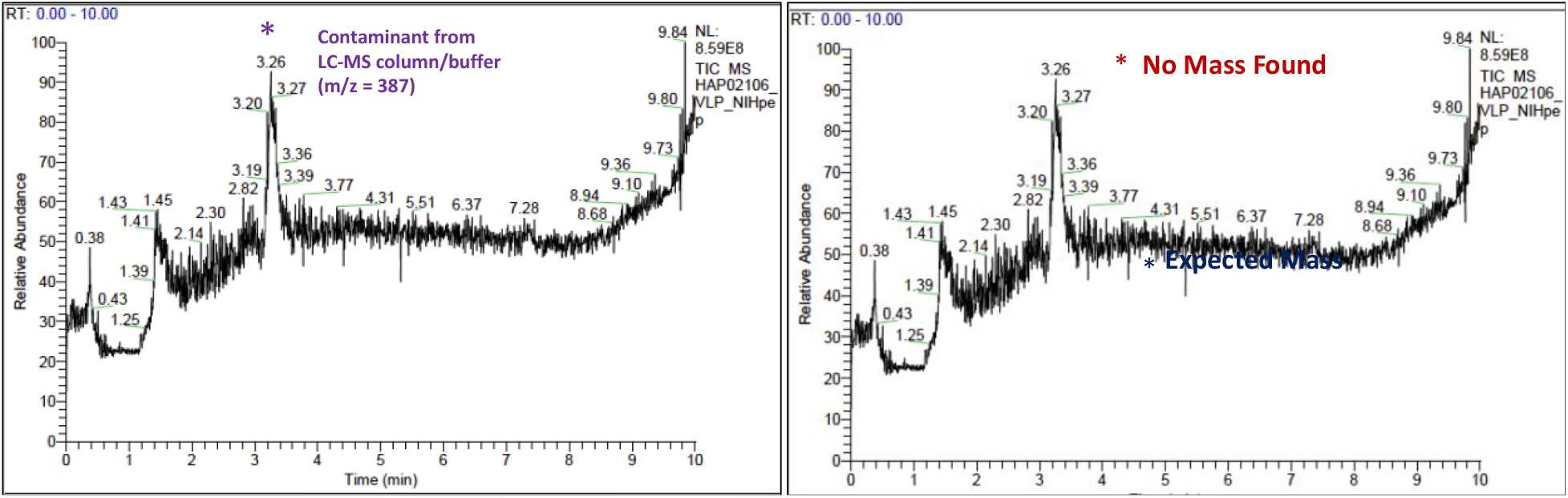
LC-MS from VLP-Open HLA-A*02:01/gTAX incubated with NIH peptide (negative control with irrelevant peptide). The peptide (YPNVNIHNF) is known to not bind A*02:01, thereby serving as a negative control. No peptide (expected mass: 1117.54) was found, as expected.

**Figure S7.**
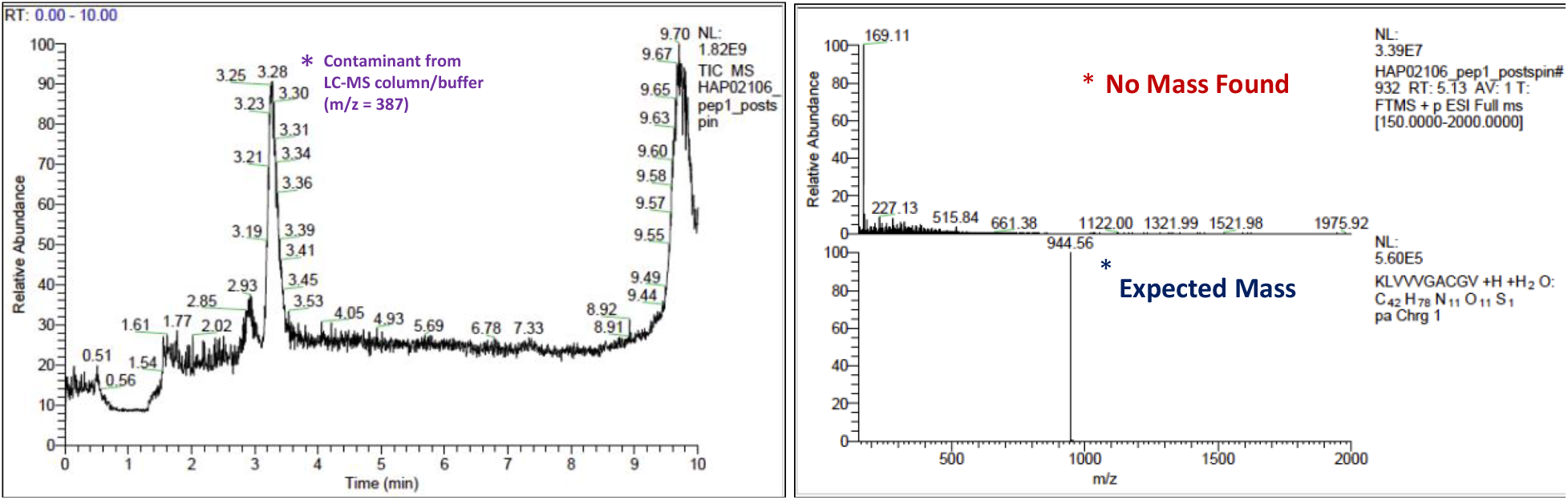
LC-MS from the sample peptide 1 (KLVVVGACGV) from TCGA library post-spinning with the 100-kDa spin column. The peptide was not found since the peptide was efficiently washed with the spin column using the same wash procedure for all other conditions. This served as a negative control, showing that only bound peptide to the VLP-Open HLA complex could be identified with mass spectrometry post-washing with the 100-kDa spin column.

**Figure S8.**
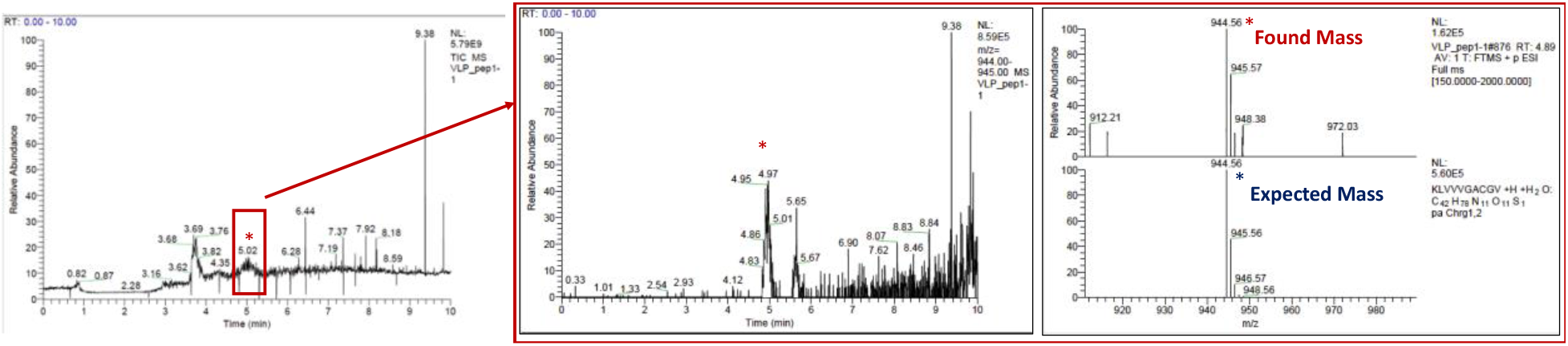
LC-MS from VLP-Open HLA-A*02:01/gTAX exchanged with peptide 1 (KLVVVGACGV) from TCGA library. The expected peptide was identified (Observed m/z: 944.57; Expected m/z: 944.56).

**Figure S9.**
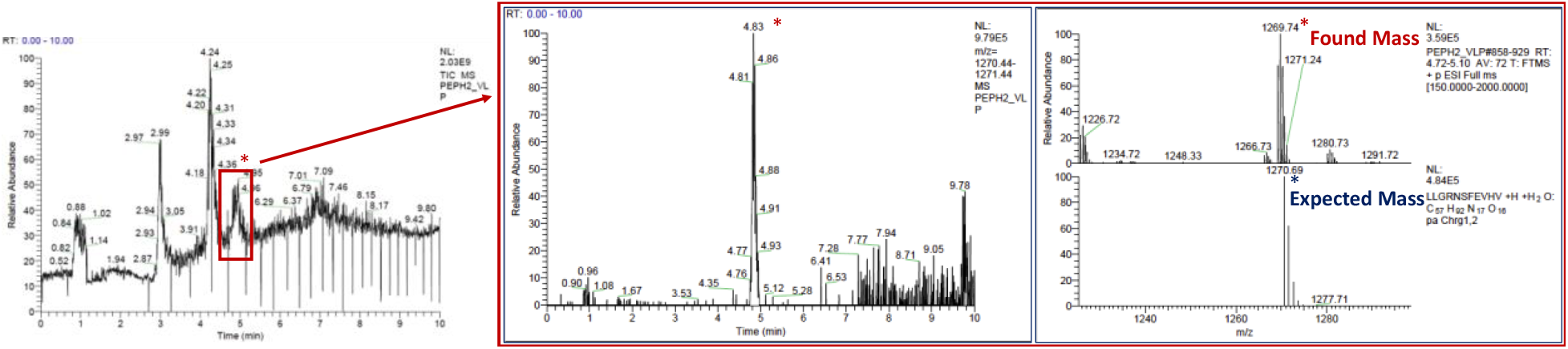
LC-MS from VLP-Open HLA-A*02:01/gTAX exchanged with peptide 2 (LLGRNSFEVHV) from TCGA library. The expected peptide was identified (Observed m/z: 1269.74; Expected m/z: 1270.69).

**Figure S10.**
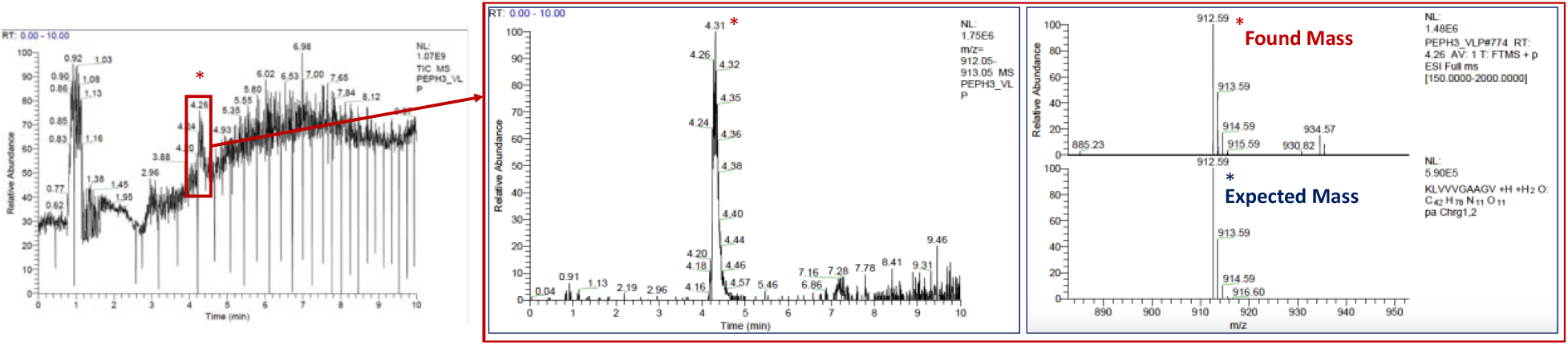
LC-MS from VLP-Open HLA-A*02:01/gTAX exchanged with peptide 3 (KLVVVGAAGV) from TCGA library. The expected peptide was identified (Observed m/z: 912.59; Expected m/z: 912.59).

**Figure S11.**
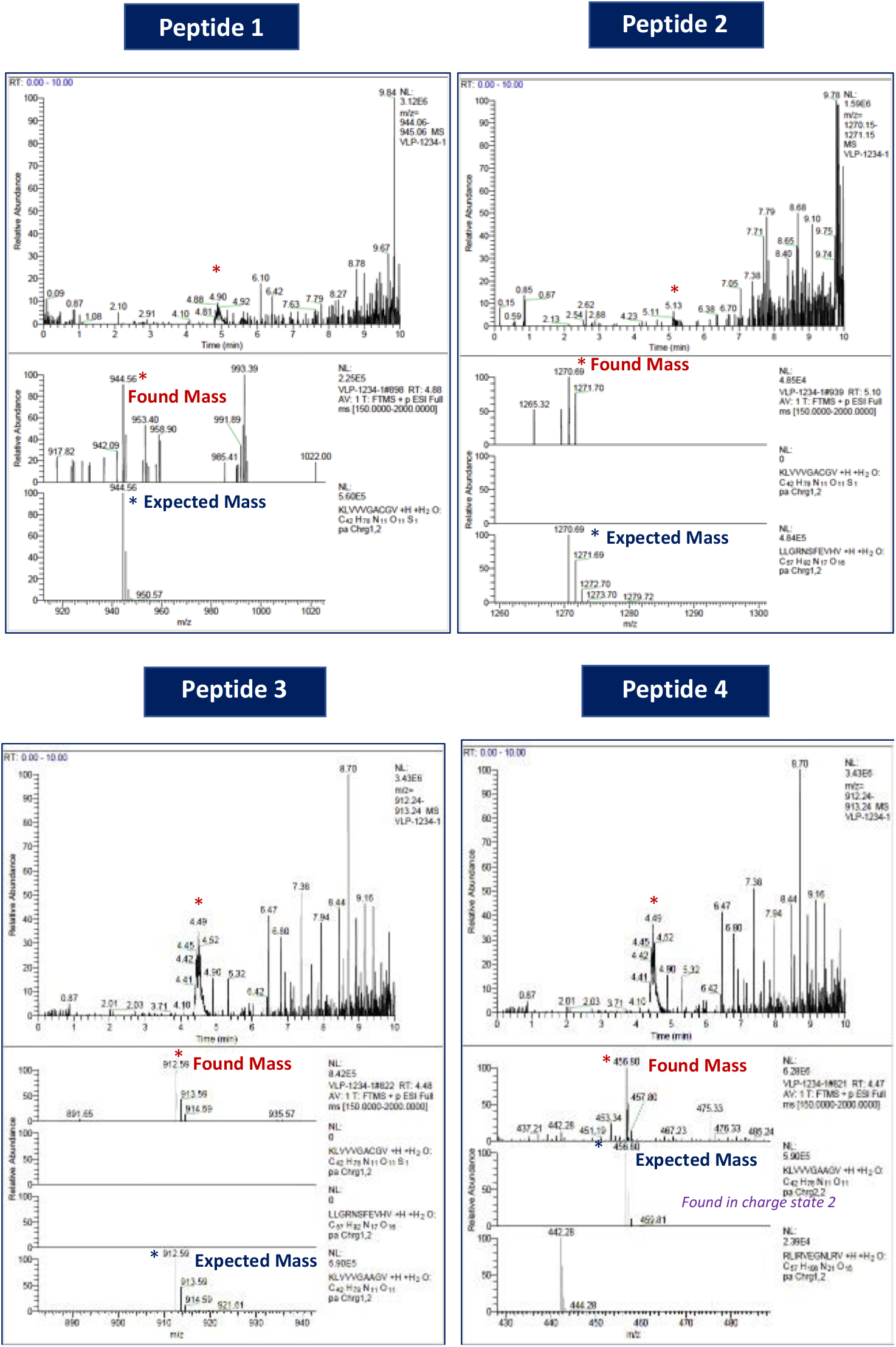
LC-MS from VLP-Open HLA-A*02:01/gTAX exchanged with a mixture of peptides 1-4 from TCGA library. All four peptides were found as indicated by the matching between the expected and the found masses.

**Figure S12.**
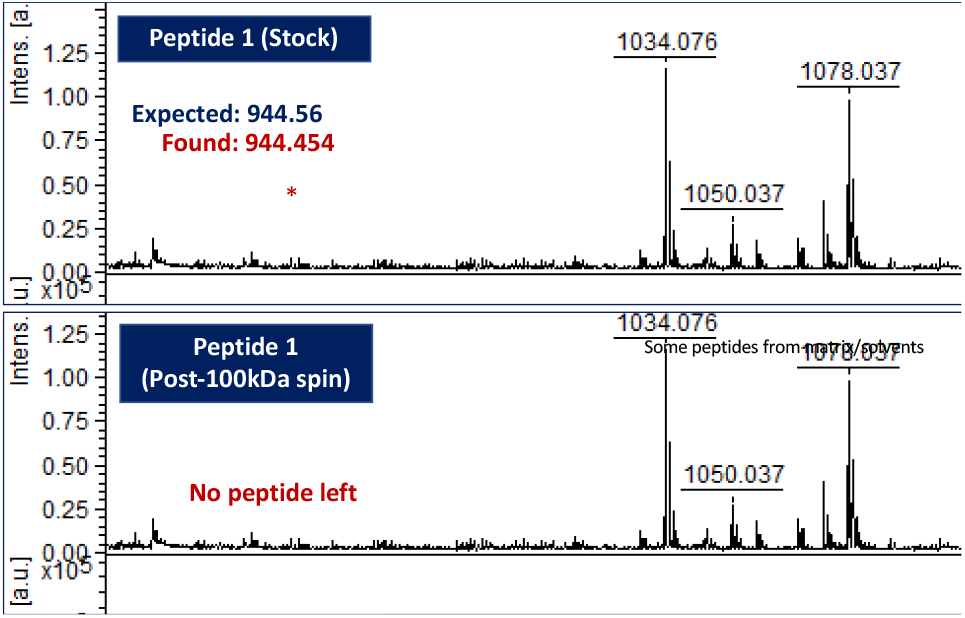
MALDI-MS from the sample peptide 1 (KLVVVGACGV) from TCGA library pre- and post-spinning with the 100-kDa spin column. *Bottom:* The peptide was not found since the peptide was efficiently washed with the spin column using the same wash procedure for all other conditions. This served as a negative control, showing that only bound peptide to the VLP-Open HLA complex could be identified with mass spectrometry post-washing with the 100-kDa spin column.

**Figure S13.**
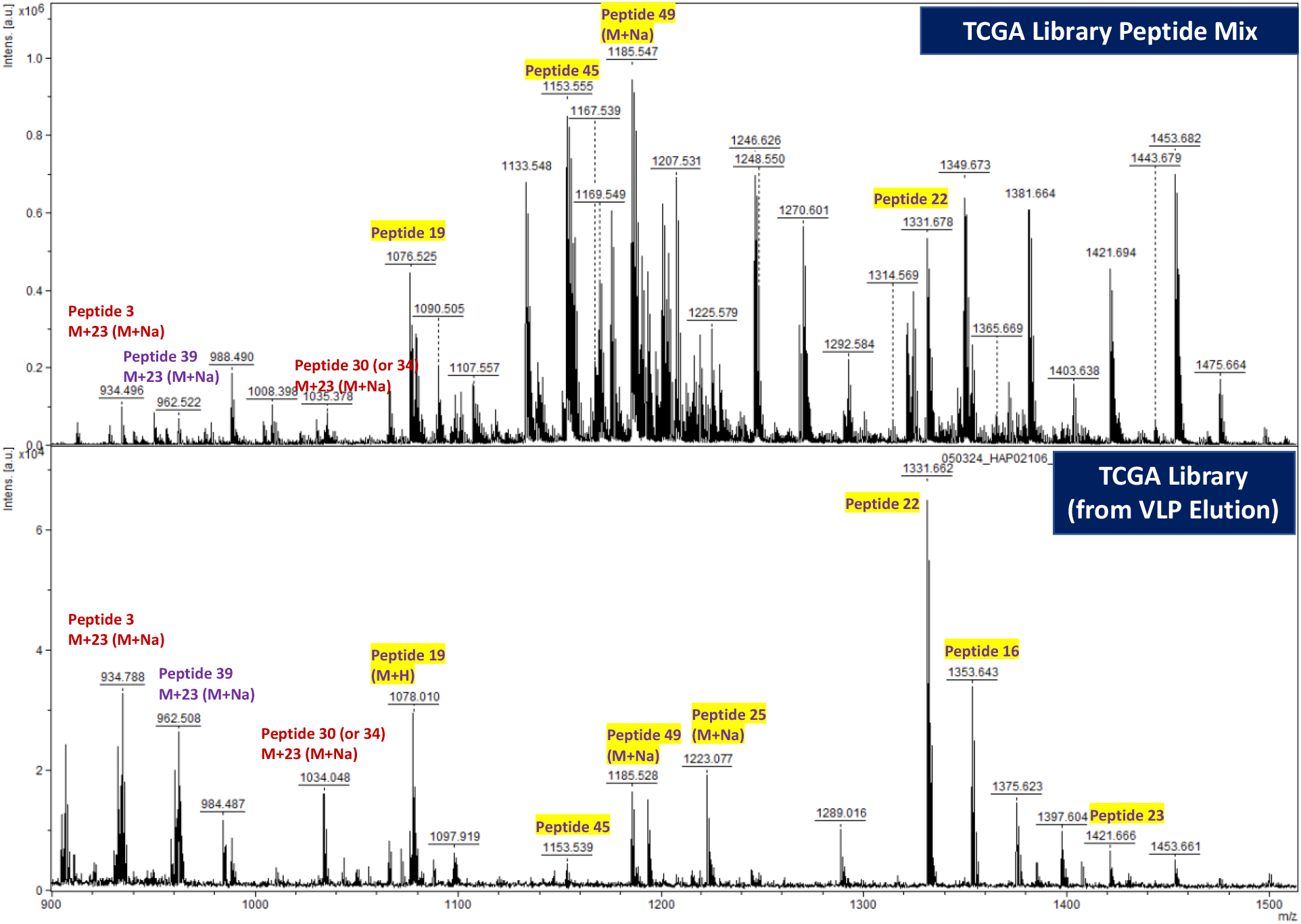
MALDI-MS from VLP-Open HLA-A*02:01/gTAX exchanged with a mixture of 50 peptides from TCGA library TCGA Library. *Top:* Mixture of 50 peptides before incubating with the VLP particles. *Bottom:* Peptides identified bound to VLP-Open HLA-A*02:01. Some found peptides are highlighted.

## 9. Fluorescent Labeling of VLP-Open HLA

SpyCatcher-mi3 (VLP) was centrifuged at 16,000 rpm for 15 minutes at 4°C to remove any potential aggregate. The particle was previously quantified using Pierce™ BCA Protein Assay. The particle was dialyzed into 50 mM borate buffer pH 8.5, to ensure that there is no sodium azide left as sodium azide can interfere with the labeling reaction. After being dialyzed overnight, the particles were concentrated using a 10kDa spin column preequilibrated with 50 mM borate buffer pH 8.5. The particle concentration was then re-quantified using Pierce™ BCA Protein Assay to ensure accurate calculation for the labeling reaction, and the VLP particles were diluted to the concentration of 2 mg/mL. NHS-fluoresceine (Thermo Scientific, Catalog #46410; molecular weight 473.4 g/mol) was equilibrated to room temperature about 20 minutes before the labeling reaction to avoid moisture. NHS-fluoresceine is reconstitute in DMSO (1 mg/100 μL). The amount of NHS-fluoresceine was calculated such that there is 15 mmol NHS-fluoresceine per 1 mmol VLP monomer. The reaction was conducted in 50 mM borate buffer 8.5 for 2 hours on ice in an amber tube shieled from light. To remove excess dye from the reaction, the Fluorescent Dye Removal Column (Thermo Scientific, Catalog #22858) was used and the protocol from the manufacturer was followed. The purified, fluorescently labeled VLP particles were then quantified using the Pierce™ BCA Protein Assay. The samples were analyzed with MALDI-MS to confirm that the labeling reaction was successful (**Figure S14**).

**Figure S14.**
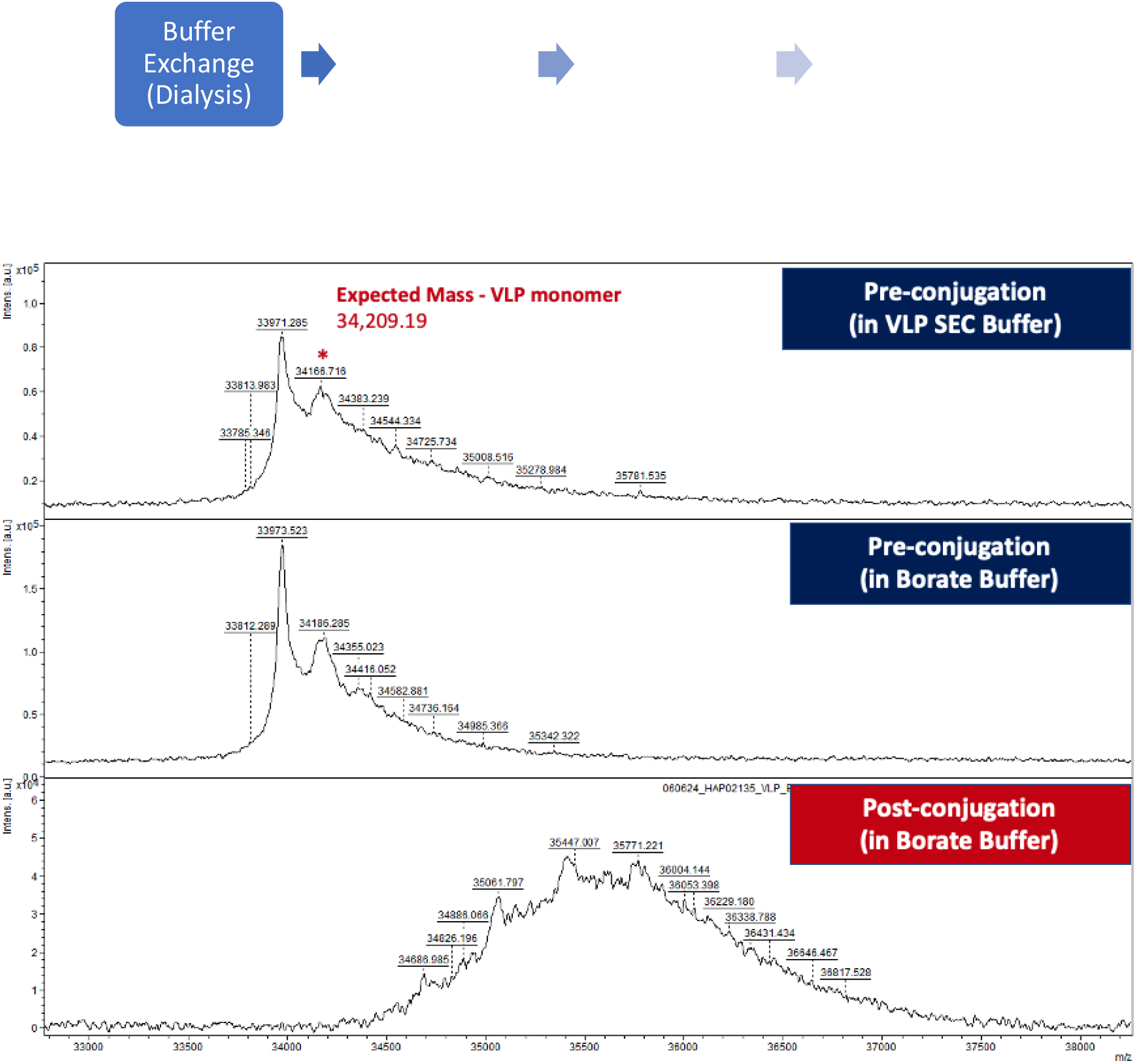
Fluorescence Label of VLP for T Cell Staining Experiment. *Top:* Labeling scheme of His6-SpyCatcher-mi3. *Bottom:* MALDI-MS of pre-labeling samples (in original SEC buffer and post-dialysis into borate buffer) and post-labeling sample. All VLP monomer (expected mass: 34,209.19) was successfully labeled as indicated by the mass shift upon labeling with NHS-fluoresceine. A mass shift corresponding to labeling of 3-5 lysine was observed.

## 10. 1G4 TCR Lentivirus and 1G4 TCR CD8+ T Cell Production

One day prior to transduction, Lenti-X 293T cells (Takara Bio) were cultured in DMEM (Gibco), 10% heat-inactivated FBS (Gibco), and Glutamax (Gibco). When cells reached a confluency between 80-90%, cells were transfected with TransIT-293 (Mirus) using pMD2.G (Addgene #12259), psPAX2 (Addgene #12260), and pSFFV-1G4. The media containing virus was collected at 24- and 48-hour post-transfection. The media was clarified by centrifuging at 500 x g for 10 minutes before being incubated with Lenti-X Concentrator (Takara Bio) for 24 hours. The virus was then pooled and concentrated 50-100 times prior to resuspension in sterile 1X PBS. The virus could be aliquoted and stored at -80°C for T cell infections. Regarding the T cells, healthy donor T cells were purchased from the Human Immunology Core at the University of Pennsylvania then CD8+ T cells were sorted out by magnetic separation prior to infection.

## 11. T Cell Staining with Fluoresceine Labeled VLP-Open HLA

CD8+ T cells 1G4 TCR, a human T cells engineered expressing the variants of a single TCR (1G4) specific for the cancer antigen NY-ESO-1, were used in these experiments. The cells were cultivated in Advanced RPMI (Gibco), supplemented with 10% heat-inactivated FBS (Gibco), 1X Glutamax (Gibco), 1X Penicillin-Streptomycin (Gibco), 10 mM HEPES (Gibco), and 10 ng/mL recombinant IL-2. In a 96-well TC-treated plate, for each well, 200 μL of T cells were cultured at 2 x 10^6^ cells/mL at 37°C and 5% CO_2_ overnight. On the next day, the cells were harvested by centrifuging the plate at 300g for 5 minutes then stained with 500 µL of 2000X diluted LIVE/DEAD Fixable Near IR Viability Stains (Invitrogen) for 10 minutes at room temperature being centrifuged at 300g for 5 minutes and washed with 500 µL FACS buffer. T cells were stained on ice with 100 µL of 1 μg/mL of either tetramers (PE-Open HLA-A*02:01/NY-ESO-1^C9V^ or PE-Open HLA-A*02:01/NY-ESO-1^W5A^ tetramers) or fluorescent-labeled VLP-Open HLA particles (VLP-Open HLA-A*02:01/NY-ESO-1^C9V^ or VLP-Open HLA-A*02:01/NY-ESO-1 ^W5A^). The samples were further stained for 20 minutes on ice with 100 µL of the diluted Brilliant Ultra Violet^TM^ 605 CD8 antibody (Biolegend, 344742) and APC anti-human TCR Vβ13.1 antibody (Biolegend, 362408). The cells were washed and centrifuged again, then fixed with 4% PFA for 30 minutes at room temperature. After being washed twice with FACS buffer, the cells were ready to be analyzed with a CytoFLEX LX flow cytometer.

For gating strategy, we identified singlets from FSC-A vs. FSC H, then identified live cells from FSC-A vs. Live-Dead IR 876-A. From this subpopulation of cells, we identified 1G4 TCR+ CD8+ by gating TCR Vβ13.1 population vs. CD8 population. The TCR Vβ13+ CD8+ cell subpopulation was gated against PE to identify cells stained with PE-tetramers (**Figure S15A**), or against FAM/fluoresceine channel to identify fluorescently labeled VLP particles (**Figure S15B**).

**Figure S15.**
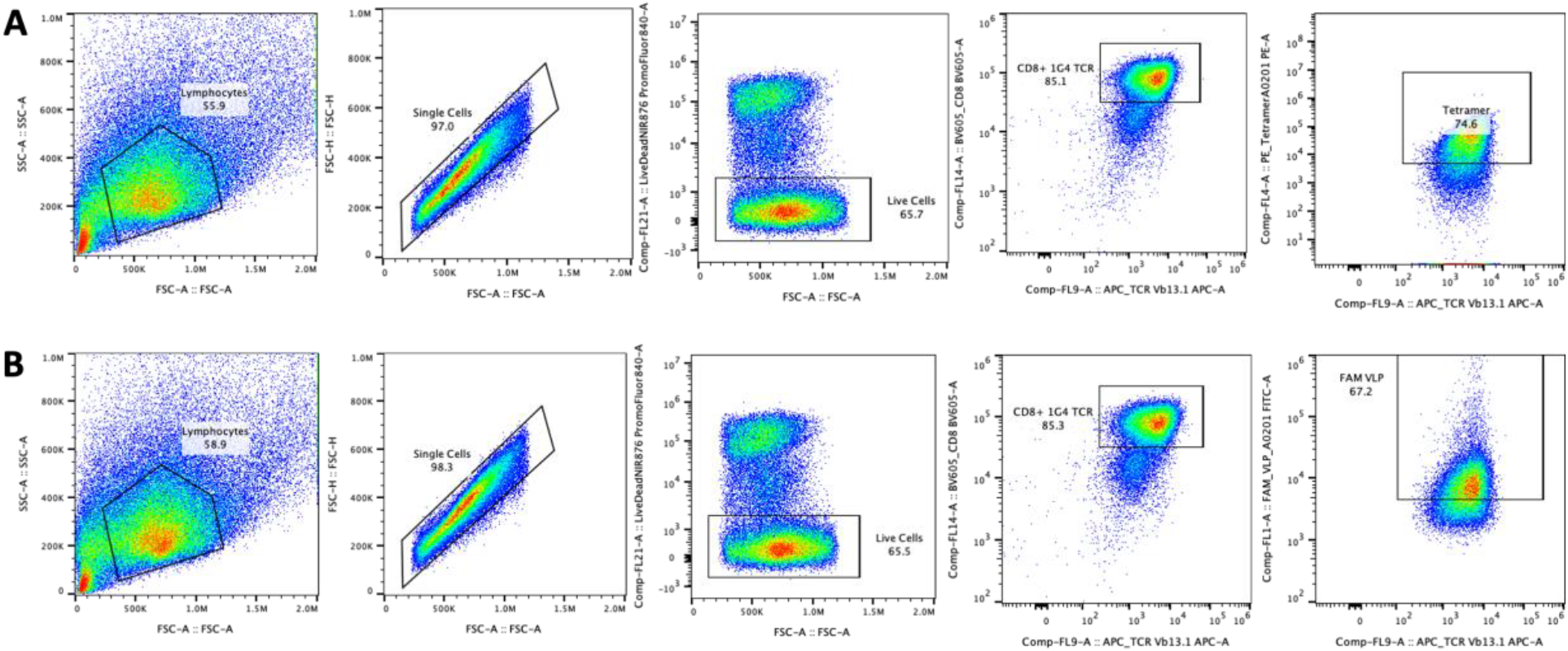
Gating Strategy for 1G4 TCR CD8+ T Cell Staining Experiment with fluorescently labeled VLP-Open HLA particles. **(A)** Gating strategy for Tetramers (controls). **(B)** Gating strategy for fluorescently labeled VLP-Open HLA particles.

## 12. T Cell Activation Assays

On Day 0 of the experiment, T cells were cultured at 1 x 10^6^ cells/mL at 37°C and 5% CO_2_ overnight. On Day 1, the cells were plated at 100,000 cells/well in a sterile U-bottomed 96-well plate; the cells could be stained with CellTrace Violet proliferation dye (Thermo) prior to stimulation according to the manufacturer’s protocol. The cells were stimulated with different concentrations of the VLP particles (VLP-Open HLA-A*02:01/NY-ESO-1^C9V^ or VLP-Open HLA-A*02:01/NY-ESO-1 ^W5A^) or at a 1:1 ratio of Dynabeads™ Human T-Activator CD3/CD28 for T Cell Expansion and Activation (Gibco) for 72 hours (3 days). A negative control with media only (no VLP and no beads) was also included. For the conditions with VLPs and the negative control, the media was also supplemented with 4 μg/mL CD28. Each condition was done in duplicates. On Day 3, harvest the cells by centrifuging the plate at 300g for 5 minutes. The cell supernatant could be saved for later analysis with ELISA. For flow cytometry analysis, the cells were stained with 500 µL of 2000X diluted LIVE/DEAD IR 876 Fixable Dead Cell Stains (Invitrogen) for 10 minutes at room temperature being centrifuged at 300g for 5 minutes and washed with 500 µL FACS buffer. The cells were centrifuged again at 300g for 5 minutes then stained with 100 µL of an antibody cocktail (Brilliant Ultra Violet™ 661 or 650 Dye* for CD69; FITC mouse anti-human for CD8, PE/Cyanine7 anti-human for CD137 or 4-1BB) in 4°C fridge for 30 mins. The cells were washed and centrifuged again, then fixed with 4% PFA for 30 minutes at room temperature. After being washed twice with FACS buffer, the cells were ready to be analyzed with a CytoFLEX LX flow cytometer.

For gating strategy, we identified singlets from FSC-A vs. FSC H, then identified live cells from FSC-A vs. Live-Dead IR 876-A. From this subpopulation of cells, we identified CD8+ by gating FSC-A vs. CD8+. The CD8+ cell subpopulation was gated against 4-1BB to identify the population expressing 4-1BB, a marker of T cell activation (**Figure S16**).

**Figure S16.**
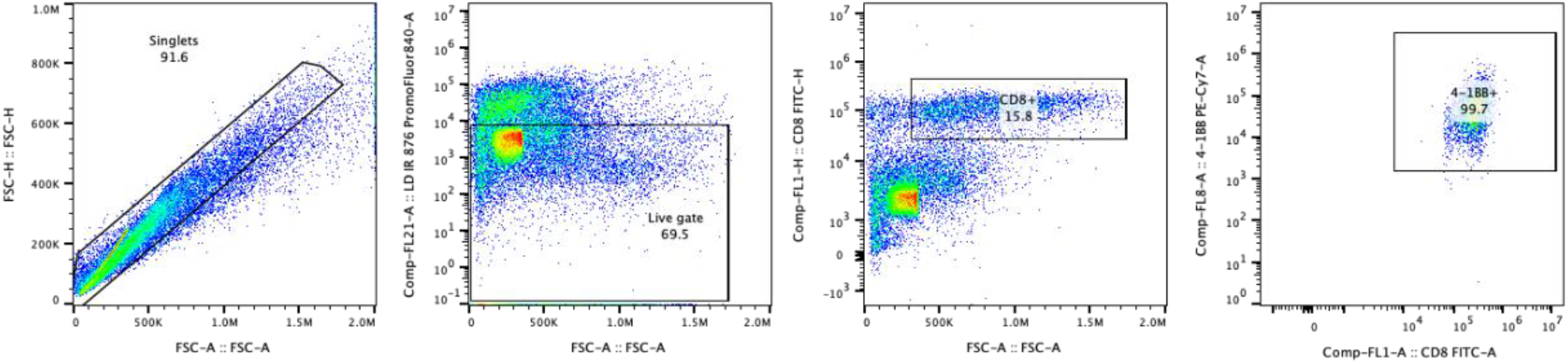
Gating strategy for T Cell activation assays with 1G4 TCR CD8+ with VLP-Open HLA-A*02:01 particles.

Data for the T cell activation assays with VLP-Open HLA-A*02:01 particles where the open HLA-A*02:01 was refolded whether with either NY-ESO-1^C9V^, NY-ESO-1^W5A^ (negative control), or gTAX (placeholder peptide) are shown in **Figure S17**. Data for the T cell activation assays with VLP-Open HLA-A*02:01/gTAX particles exchanged with either NY-ESO-1^C9V^ or NY-ESO-1^W5A^ are shown in **Figure S18**. Data for the MFI of Eomes of CD8+ cells are shown in **Figure S21**.

**Figure S17.**
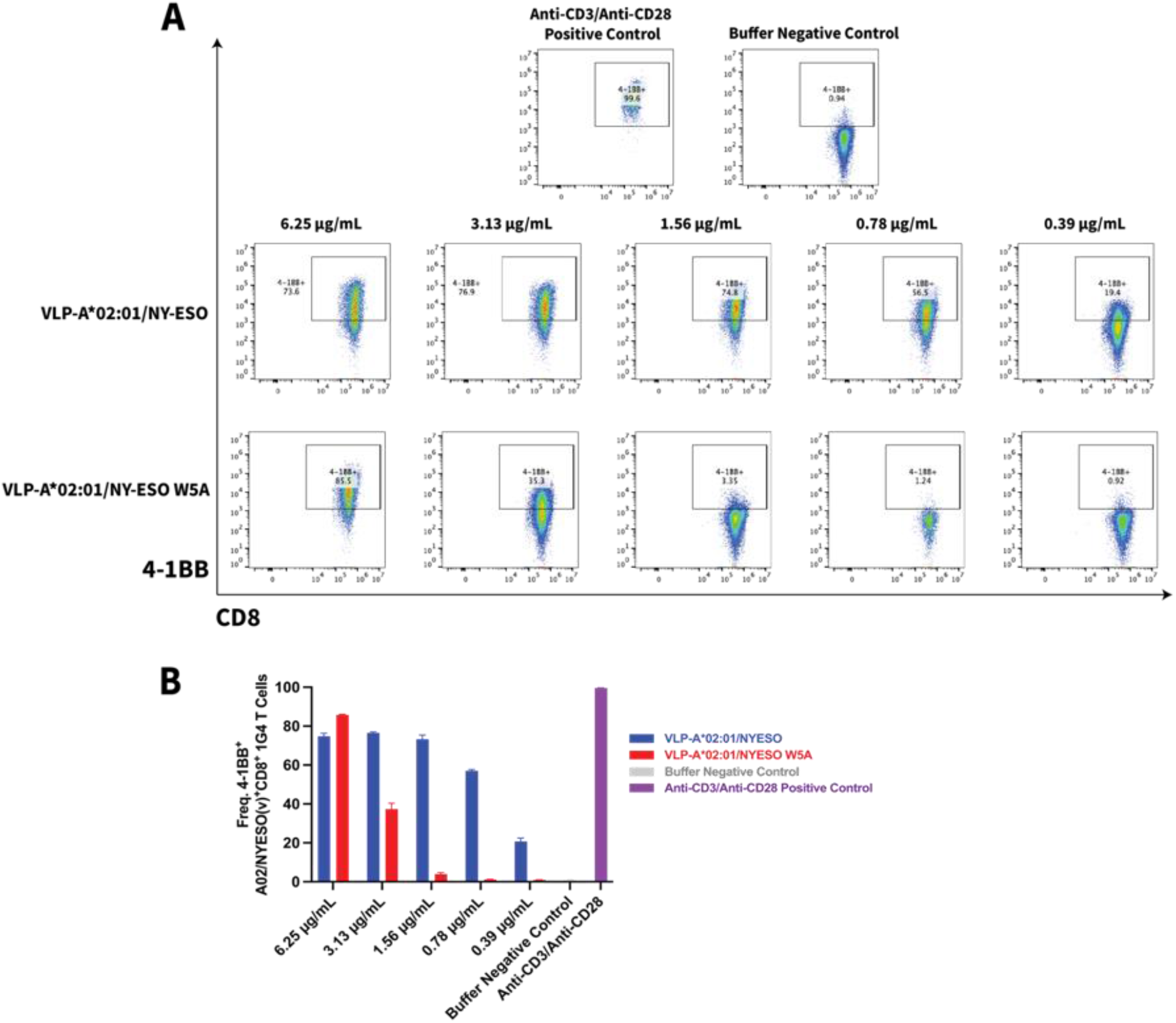
Exploring the Optimal Working Concentration Range of VLP-Open HLA-A*02:01/NY-ESO-1^C9V^ for Specific T cell Activation in a Cognate Antigen Dependent Manner. 1G4 CD8+ T cells were stimulated with different concentrations of VLP conjugated with Open HLA refolded with peptides: VLP-Open HLA-A*02:01/NY-ESO-1^C9V^, VLP-Open HLA-A*02:01/NY-ESO-1^W5A^, media only (buffer negative control), or anti-CD3/anti-CD28 beads (positive control) for 3 days before flow cytometry analysis. **(A)** Representative flow cytometry data of 4-1BB+ CD8+ cell populations across different conditions. **(B)** Frequency of 4-1BB+ cells in different conditions. The average of at least 2 technical replicates is shown.

**Figure S18.**
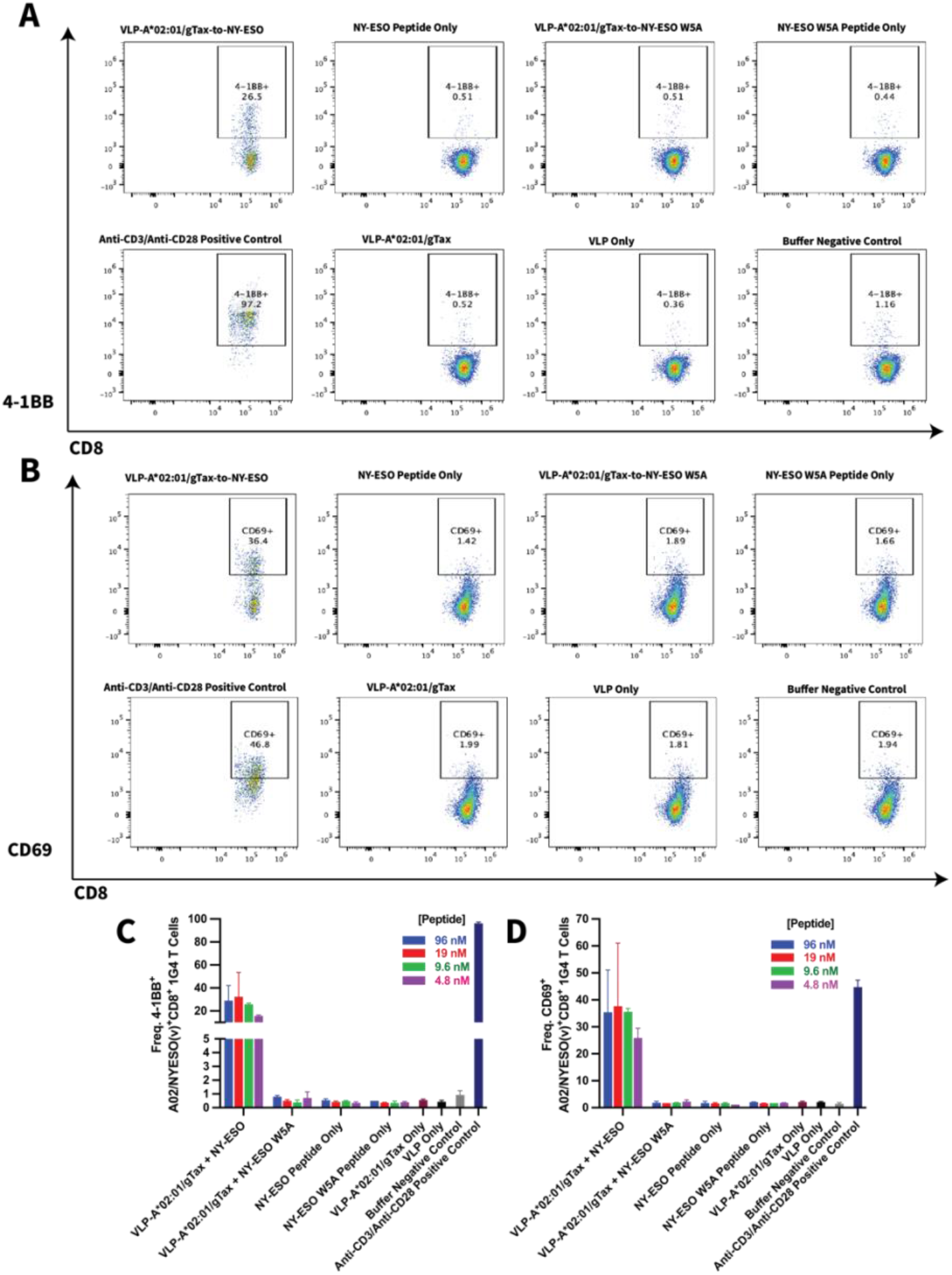
VLP-Open HLA-A*02:01/gTax exchanged with NY-ESO-1^C9V^ can trigger T cell activation in a cognate antigen dependent manner. 1G4 CD8+ T cells were stimulated with VLP-Open HLA-A*02:01/gTax exchanged with different concentrations of NY-ESO-1^C9V^ or NY-ESO-1^W5A^, VLP-HLA-A*02:01/gTax only (no exchange), VLP only, peptide only (NY-ESO-1^C9V^ or NY-ESO-1^W5A^), media only (buffer negative control), or anti-CD3/anti-CD28 beads (positive control) for 3 days before flow cytometry analysis. **(A)** Representative flow cytometry data of 4-1BB+ CD8+ cell populations across different conditions. **(B)** Representative flow cytometry data of CD69+ CD8+ cell populations across different conditions. **(C)** Frequency of 4-1BB+ cells in different conditions. **(D)** Frequency of CD69+ cells in different conditions. The average of at least two technical replicates is shown.

## 13. ELISA

The cell supernatant from T cell activation assays were stored in -20°C freezer until analysis by ELISA. The samples were diluted 2X-3X for ELISA analysis with DuoSet ELISA: Human IFN-γ (R&D-Biotechne; DY285B) and DuoSet Ancillary Reagent Kit (R&D – Biotechne; DY008) according to the manufacturer’s protocol. In brief, a flat-bottom polystyrene microplate was coated with 2.00 μg/mL capture antibody overnight before being washed and blocked with the Reagent Diluent containing 1% BSA. In addition to the samples, a standard curve with recombinant human IFN-γ was prepared in Reagent Diluent was prepared, and they were added to the plate with an incubation time of 2 hours at room temperature. The plate was washed with the Wash Buffer before adding Detection Antibody (biotinylated mouse anti-human IFN-γ) at 200 ng/mL for another 2-hour incubation at room temperature. The plate was washed and 1X Streptavidin-HRP B was added to the plate for 20 minutes incubation at room temperature. The plate was washed again before TMB ELISA Substrate was added to each well for a 20-minute incubation at room temperature. A stop solution (2N sulfuric acid) was added to each well to stop the reaction. The OD at 450 nm (correction at 540 nm or 570 nm) was collected using a SpectraMax iD5 plate reader. ELISA analysis of IFN-γ from the cell supernatant of T cell activation assays with VLP-Open HLA-A*02:01 particles where the open HLA-A*02:01 was refolded whether with either NY-ESO-1^C9V^, NY-ESO-1^W5A^ (negative control), or gTAX (placeholder peptide) is shown in **Figure S19**. ELISA analysis of IFN-γ from the cell supernatant of the T cell activation assays with VLP-Open HLA-A*02:01/gTAX particles exchanged with either NY-ESO-1^C9V^ or NY-ESO-1^W5A^ are shown in **Figure S20**.

**Figure S19.**
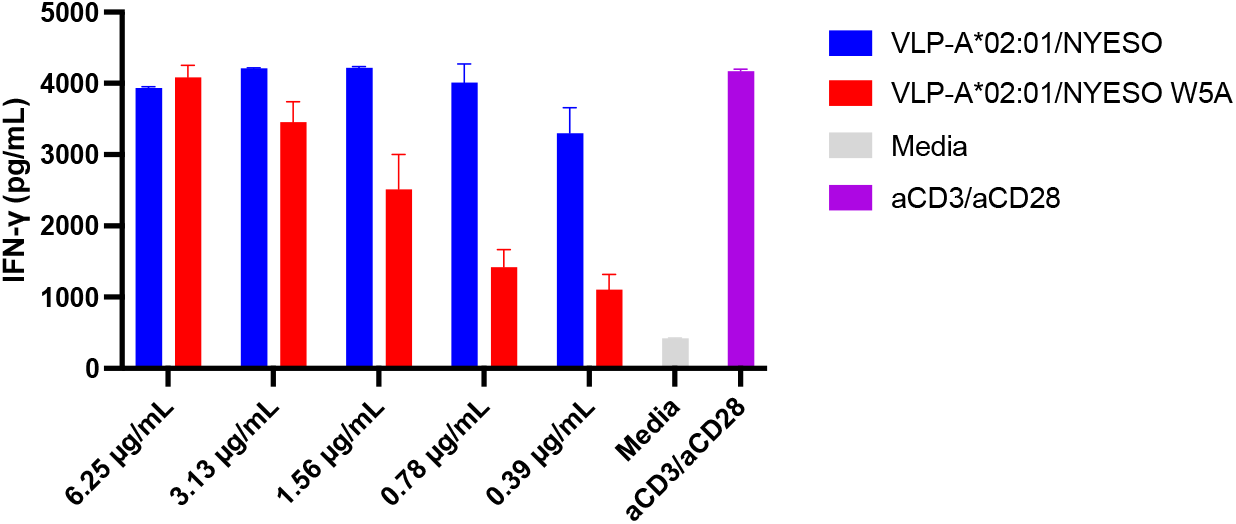
ELISA analysis of IFN-γ from the cell supernatant of T cell activation assays with VLP-Open HLA-A*02:01 particles where the open HLA-A*02:01 was refolded whether with either NY-ESO-1^C9V^, NY-ESO-1^W5A^ (negative control), or gTAX (placeholder peptide). The condition with media or buffer only served as the negative control. The condition with anti-CD3 anti-CD28 beads served as the positive control for activation. The average of at least 2 technical replicates is shown.

**Figure S20.**
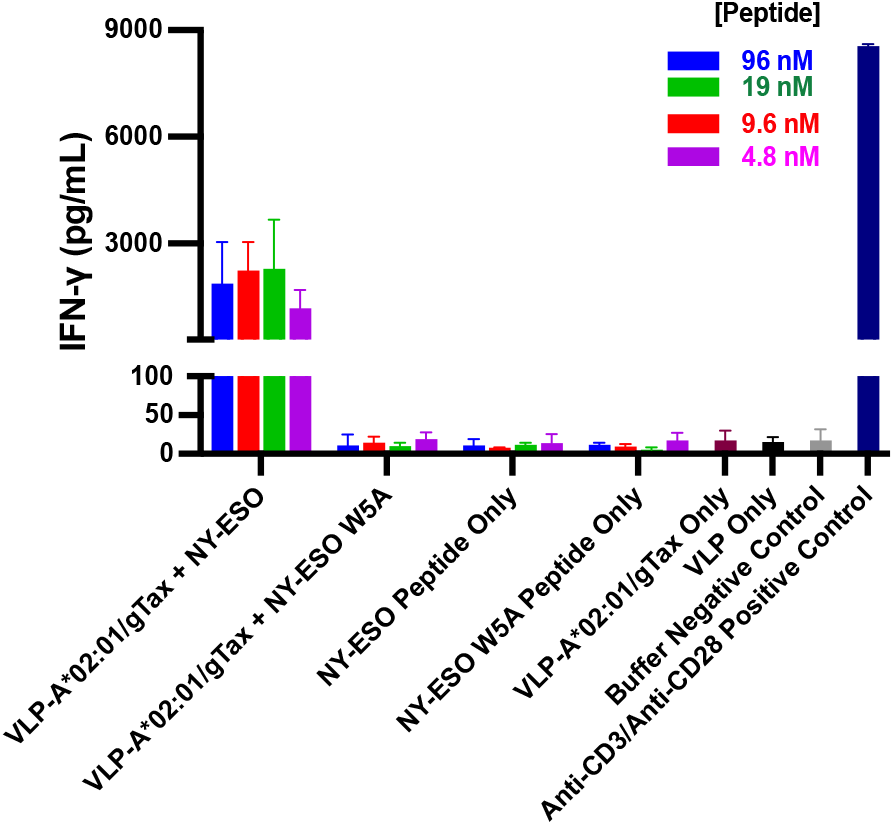
ELISA analysis of IFN-γ from the cell supernatant of the T cell activation assays with VLP-Open HLA-A*02:01/gTAX particles exchanged with either NY-ESO-1^C9V^ or NY-ESO-1^W5A^. The conditions with peptide only, VLP-Open HLA-A*02:01/gTAX only, VLP only, and media or buffer only served as the negative control. The condition with anti-CD3 anti-CD28 beads served as the positive control for activation. The average of at least 2 technical replicates is shown.

**Figure S21.**
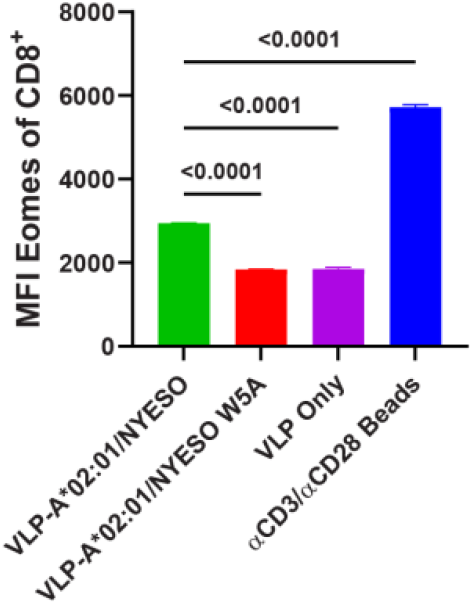
1G4 T cells were stimulated with VLP-Open HLA-A*02:01/NY-ESO-1^C9V^, VLP-Open HLA-A*02:01/NY-ESO-1^W5A^, VLP only (negative control), or anti-CD3/anti-CD28 beads. VLP particles were added at a concentration 0.78 μg/mL and incubated for 3 days. MFI of Eomes among CD8+ cells is shown.

